# Sugarcane yellow leaf virus impairs the transcriptomic defense responses of sugarcane to its new aphid vector *Melanaphis sorghi*

**DOI:** 10.1101/2024.10.17.618922

**Authors:** Ricardo José Gonzaga Pimenta, Alexandre Hild Aono, Danilo Augusto Sforça, Maria Natália Guindalini Melloni, Marcos Cesar Gonçalves, Luciana Rossini Pinto, Anete Pereira de Souza

## Abstract

Sugarcane (*Saccharum* spp.) is a preeminent sugar and bioenergy crop and has great economic importance in tropical countries. A major disease affecting this crop is yellow leaf disease, which is caused by sugarcane yellow leaf virus (SCYLV, *Polerovirus SCYLV*, *Solemoviridae*). The sugarcane aphid *Melanaphis sacchari* is considered the main vector of SCYLV, and the closely related sorghum aphid *Melanaphis sorghi,* which has recently emerged as a pest of great relevance in sorghum, has also been suspected as a vector. Genetic resistance is an important resource for preventing yield losses caused by SCYLV and its vectors, but knowledge on the underlying molecular mechanisms is lacking. Therefore, the present work was aimed at investigating the transcriptomic responses of sugarcane to SCYLV and *M. sorghi*, which was reported to transmit this virus for the first time herein. Two sugarcane cultivars, one susceptible and one tolerant to SCYLV, were fed upon by aviruliferous and viruliferous aphids. The transcriptome of the plants was assessed by RNA-Seq using differential gene expression analyses and a gene coexpression network. The susceptible cultivar showed an incipient reaction to both *M. sorghi* and SCYLV, with very few differentially expressed genes (DEGs) identified in comparison with aphid-free plants. The response of the tolerant cultivar to aviruliferous *M. sorghi* involved pathways typically associated with defense against herbivory, which were also enriched in coexpression network modules in which DEGs were overrepresented. Some of these genes were hubs in their respective modules, indicating they are potential key regulators of the defense responses. However, these responses were diminished when viruliferous aphids were used, and other processes linked to infection with SCYLV were altered. These results indicated that SCYLV could affect sugarcane defense responses to its vector, similar to other viruses of the same genus. Some possible implications for the epidemiology and impact of SCYLV and *M. sorghi* are discussed.

## 1. Background

Sugarcane (*Saccharum* spp.) is a major sugar and energy crop in tropical regions of the world, accounting for nearly 80% of the global production of sugar (ISO 2023). The yield of this crop is affected by several diseases, including yellow leaf disease (YLD), which is recognized mainly by the yellowing of abaxial leaf midribs of mature plants (Vega et al. 1997; ElSayed et al. 2015). This disease is caused by sugarcane yellow leaf virus (SCYLV, *Polerovirus SCYLV*), a positive-sense ssRNA virus belonging to the genus *Polerovirus* of the *Solemoviridae* family (Sõmera et al. 2021) present in all sugarcane-growing regions of the world (Komor et al. 2010). This virus is known to affect photosynthesis efficiency as well as the metabolism and transport of carbohydrates (Gonçalves et al. 2005; Lehrer et al. 2010; Viswanathan et al. 2014). This impairs growth and leads to substantial reductions in sugar productivity and tonnage, which reflect yield losses of up to 37% (Rassaby et al. 2003; Vasconcelos et al. 2009; Viswanathan et al. 2014).

SCYLV is transmitted in a persistent, circulative and nonpropagative manner by several aphid species, among which the sugarcane aphid *Melanaphis sacchari* (Zehntner, 1897) is considered the most relevant (Scagliusi and Lockhart 2000). *M. sacchari* is capable of reproducing by apomictic parthenogenesis and is widely distributed throughout the world (Singh et al. 2004). In sugarcane, economic losses caused by *M. sacchari* are usually not concerning, and the species is mostly regarded as an important vector of viruses (Hall 1987; White et al. 2001). In the last decade, a ‘superclone’ of a closely related species, the sorghum aphid *Melanaphis sorghi* (Theobald, 1904), has emerged as a major pest of sorghum (*Sorghum bicolor*) in the Americas (Nibouche et al. 2018, 2021). Although the transmission of SCYLV by *M. sorghi* has not been reported thus far, SCYLV has been detected in *S. bicolor* (Wei et al. 2016; ElSayed et al. 2018).

While the physiological consequences of infection by SCYLV and its impact on yield have been widely assessed in sugarcane, knowledge on the molecular mechanisms that underlie the response to this virus is still lacking. Genetic mapping studies targeting SCYLV resistance have been performed but are limited by the highly complex sugarcane genome, phenotyping constraints, and the quantitative nature of this trait (Debibakas et al. 2014; Pimenta et al. 2021). The ineffectiveness of the transmission of SCYLV by mechanical methods (Scagliusi and Lockhart 2000) has further restricted research on this virus, as it implies that data on the resistance to SCYLV and to its vectors are frequently intertwined.

For instance, a marker associated with resistance to SCLYV identified by association mapping was found to be linked to a lipoxygenase gene associated with aphid resistance (Debibakas et al. 2014). Sugarcane resistance to aphids, which could help prevent infection by SCYLV, has been assessed at the phenotypic level (e.g., Akbar et al. 2011; Fartek et al. 2014; Bertasello et al. 2021), but its genetic basis has not been investigated in this crop.

With the decreasing cost of high-throughput next-generation sequencing technologies, RNA sequencing (RNA-Seq) has become a common method for investigating the mechanisms underlying biological phenomena. Research on sugarcane has contributed to the understanding of plant responses to pests and several pathogens, including viruses (Ling et al. 2018; Akbar et al. 2020). This type of analysis has not yet been undertaken for the feeding of sucking insects and has been performed for SCYLV infection only in a susceptible cultivar infested with aviruliferous and viruliferous *M. sacchari* (Shabbir et al. 2022). Assessing the responses of genotypes with contrasting responses to SCYLV, as well as those of aphid-free plants, can help elucidate the molecular mechanisms underlying resistance to both aphids and viruses, as well as their interactions.

In this study, the ability of *M. sorghi* to transmit SCYLV in sugarcane was demonstrated, with transmission rates reaching 100% after 48 hours of feeding. Subsequently, the transcriptomic responses of sugarcane to SCYLV and *M. sorghi* were investigated using RNA-Seq. Two sugarcane hybrid cultivars, one susceptible and one tolerant to SCYLV, were exposed to either aviruliferous or viruliferous aphids. Comparing the transcriptomes of these plants with those of aphid-free controls and building a weighted gene coexpression network provided insights into the early molecular responses of sugarcane to SCYLV and *M. sorghi*.

## 2. Results

### 2.1. Viral isolate and aphid species identification

The complete genome of the SCYLV isolate infecting the source of inoculum, spanning 5,903 nucleotides, was obtained by Sanger sequencing and pairwise-aligned with publicly available genomes of 101 SCYLV isolates. The genome of this new isolate, named BRA-SP1, was found to be most closely related to that of BR-06-Leaf, which originated from a sugarcane sample collected in 2018 in Brazil. Forty-one polymorphisms were detected between the genomes of BRA-SP1 and BR-06-Leaf, which presented 99.3% similarity.

Recombination analyses indicated that thirteen isolates included in the alignment were potential recombinants (Table S1), and these were thus excluded from the subsequent phylogenetic analysis to avoid confounding effects. A maximum likelihood tree was constructed with the remaining 89 aligned sequences using a general time reversible model with gamma-distributed invariant sites. The results indicated the existence of three well-supported major SCYLV clades, including all the isolates from Mauritius and Reunion (clade REU), a few isolates from Asia, the Americas and Papua New Guinea (clade CUB), and isolates from diverse locations (clade BRA). Isolate BRA-SP1 was positioned in clade BRA, close to other samples originating from Brazil (Figure 1).

**Figure 1.**
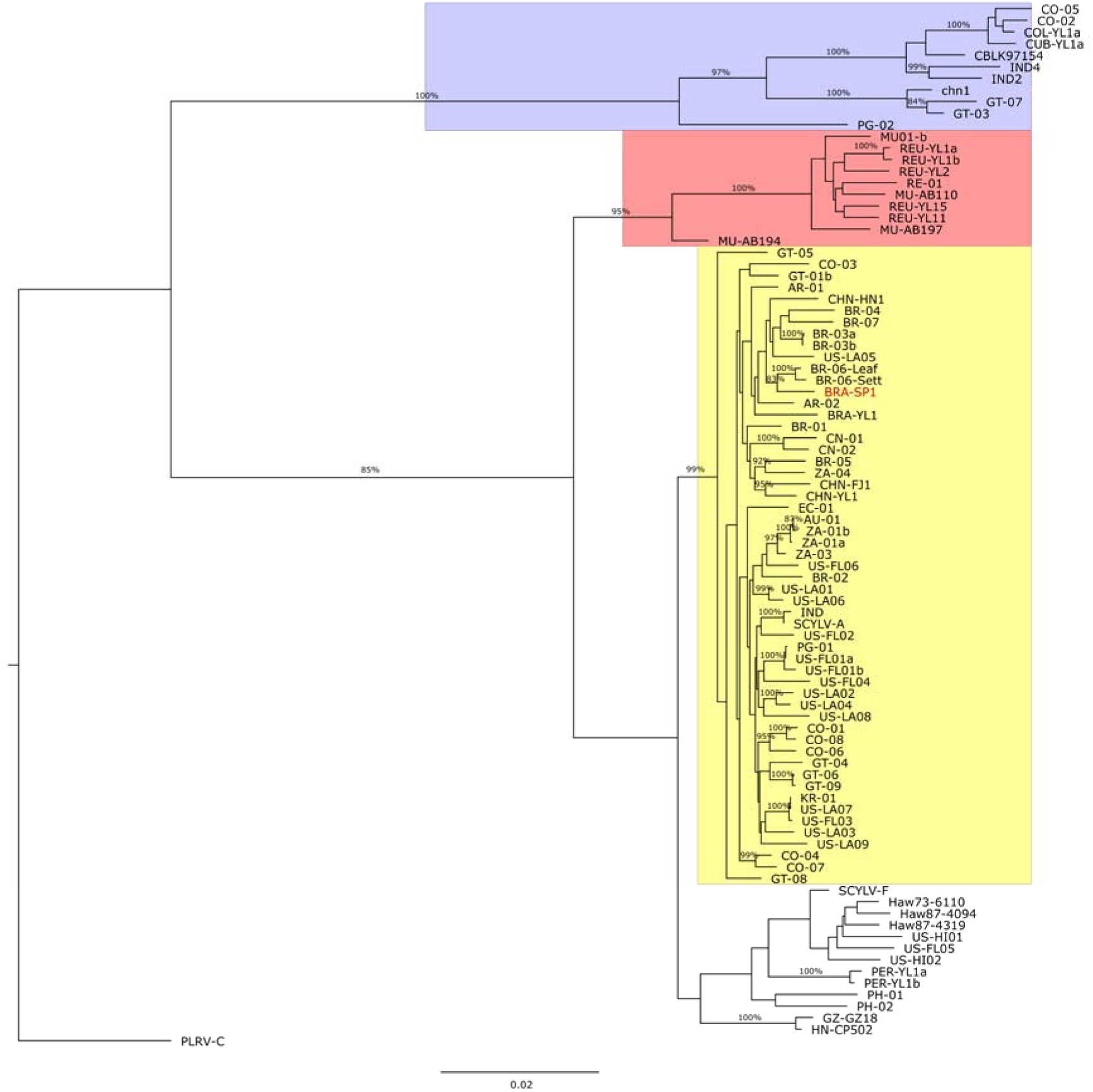
Maximum likelihood phylogenetic tree constructed using complete genome sequences of 89 sugarcane yellow leaf virus isolates. Numbers on the branches represent bootstrap values, for which a cutoff of 80% was used. The scale bar signifies a genetic distance of 0.02 nucleotide substitutions per site. Three major clades were identified: CUB (blue), REU (red) and BRA (yellow).

A colony of aviruliferous *Melanaphis* sp. aphids collected from a sugarcane field was established, and a fragment of the *elongation factor 1-alpha* (*EF1-*α) gene harboring five polymorphic sites was sequenced from a pool of five insects. The nucleotides observed at positions 296, 478, 637, 663, and 735 were Y, T, A, G, and C, respectively. These results match the *EF1-*α haplotype “H2c” described by Nibouche et al. (2021), and thus the colony was identified as *M. sorghi*.

### 2.2. Inoculation experiment

Plants of the sugarcane cultivars SP71-6163 (SCYLV-susceptible) and IACSP95-5000 (SCYLV-tolerant) were exposed to *M. sorghi* aphids for varying periods to assess viral transmission. Reverse transcription followed by quantitative PCR (RT[qPCR) confirmed that SCYLV was not present in any plants exposed to aviruliferous aphids or plants not exposed to aphids, which were used as controls. The number and percentage of plants in which SCYLV was detected at different inoculation access periods (IAPs) are summarized in Table S2. The virus was detected in SP71-6163 after as little as four hours of exposure to *M. sorghi*, showing that this aphid is capable of vectoring SCYLV in sugarcane under controlled conditions. The virus was detected in 100% of the plants of both cultivars after only 48 hours of exposure to *M. sorghi*; thus, this IAP was chosen for subsequent RNA-Seq analysis. At this time point, the mean SCYLV relative quantification of SP71-6163 (0.0013) was significantly (*p* = 0.03) greater than that of IACSP95-5000 (0.0003).

### 2.3. Transcriptome reference assembly and quantification

The sequencing of the 18 RNA-Seq libraries yielded more than a billion paired-end reads; over 95% of these (961.2 million) could be attributed to the libraries, resulting in an average of 53.4 million reads per sample. After trimming, all reads presented an average Phred score ≥ 24, and 99.7% of them presented a Phred score ≥ 30 (Table S3).

In conjunction with these data, trimmed RNA-Seq data from leaf samples of twelve diverse *Saccharum* genotypes (Correr et al. 2020) were used to construct a new reference sugarcane transcriptome. Filtered reads were aligned to the allele-resolved genomes of the two main ancestral *Saccharum* species: *Saccharum spontaneum* and *Saccharum officinarum*. The numbers of assembled transcripts for the *S. spontaneum* haplotypes were (A) 33,969, (B) 33,913, (C) 33,576 and (D) 33,672. For *S. officinarum*, the quantities were as follows: (A) 34,993, (B) 34,081, (C) 33,832, (D) 32,513, (E) 31,720, (F) 30,280, (G) 29,124 and (H) 25,926. Reduction of redundancies in the two references using CD-HIT resulted in 95,533 and 141,626 transcripts for *S. spontaneum* and *S. officinarum*, respectively. These two sets were then combined, yielding a final reference containing 206,426 transcripts with an N50 of 2,097 bases. Searches using BUSCO showed that this reference included 99.3 and 98.4% of the conserved orthologs of eukaryotes and green plants, respectively (Table S4). A total of 43,291 of these transcripts, hereafter referred to as “genes”, could be annotated with Trinotate.

On average, 89.4% of the reads from each RNA-Seq library generated in the present study were mapped to this reference (Table S3). The first two gene filtering steps based on count per million (CPM) values resulted in the retention of 85,685 expressed genes. A multidimensional scaling analysis performed with these data is shown in Figure S1, where the proximity between replicates of each genotype and biological condition is observed.

### 2.4. Differential gene expression analyses

After performing t tests, sets of genes to be excluded in each condition were obtained and combined according to the comparisons to be carried out in the differential gene expression (DGE) analyses. The quantities of genes excluded in each condition and in the union of sets, as well as the number of genes retained for each comparison, can be found in Table S5. DGE analyses were performed for nine different comparisons. The number of differentially expressed genes (DEGs) identified in each group can be seen in Table 1, which distinguishes upregulated (log_2_(fold change) (log_2_(FC)) > 1) and downregulated (log_2_(FC) < 1) genes, and in Figure 2, which shows intersections between DEG sets. Additionally, Table S6 presents a complete list of DEGs and associated information. The expression patterns of eight DEGs were validated through RT□qPCR using *glyceraldehyde-3-phosphate dehydrogenase* (*GAPDH*) and Cluster_58111 as endogenous controls. Overall, strong and significant linear correlations between the CPM and ΔCq values were observed, with R^2^ values ranging from 0.56 to 0.92 (Figure S2).

**Figure 2.**
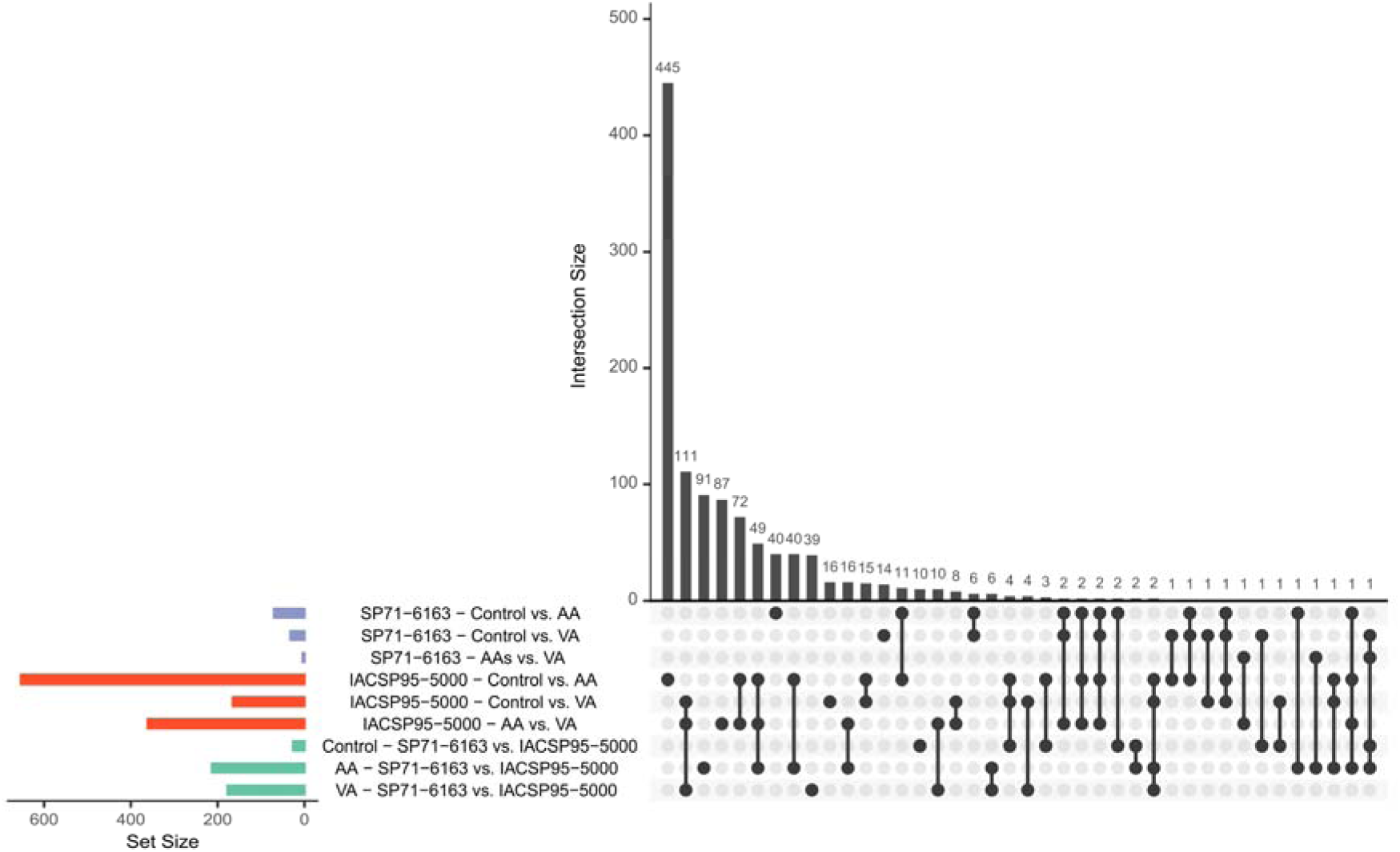
Upset plot displaying intersections between differentially expressed gene sets in SP71-6163 and IACSP95-5000.

**Table 1.**
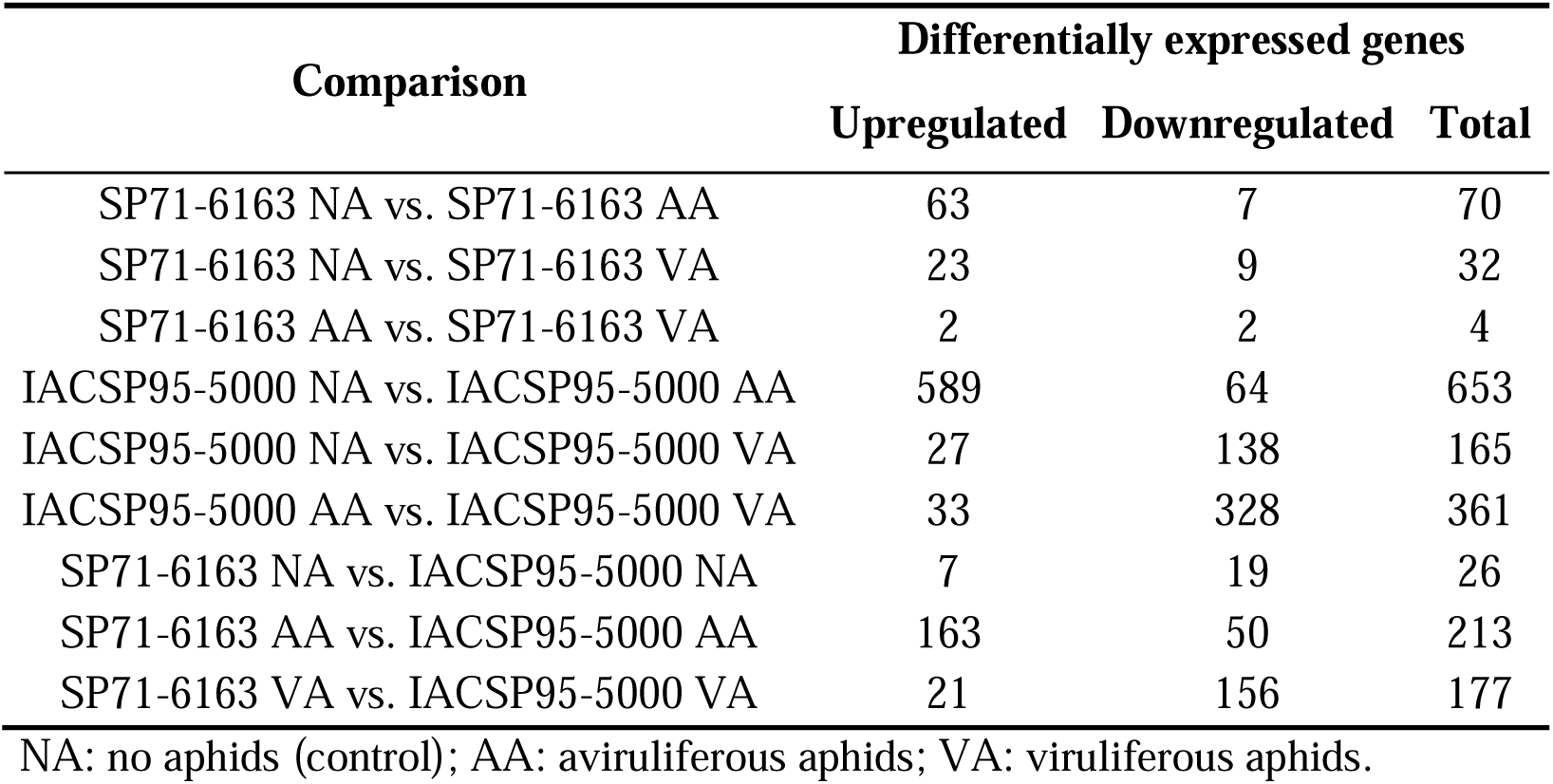
Number of up- and downregulated differentially expressed genes in each comparison performed.

| Comparison | Differentially expressed genes |  |  |
| --- | --- | --- | --- |
|  | Upregulated | Downregulated | Total |
| SP71-6163 NA vs. SP71-6163 AA | 63 | 7 | 70 |
| SP71-6163 NA vs. SP71-6163 VA | 23 | 9 | 32 |
| SP71-6163 AA vs. SP71-6163 VA | 2 | 2 | 4 |
| IACSP95-5000 NA vs. IACSP95-5000 AA | 589 | 64 | 653 |
| IACSP95-5000 NA vs. IACSP95-5000 VA | 27 | 138 | 165 |
| IACSP95-5000 AA vs. IACSP95-5000 VA | 33 | 328 | 361 |
| SP71-6163 NA vs. IACSP95-5000 NA | 7 | 19 | 26 |
| SP71-6163 AA vs. IACSP95-5000 AA | 163 | 50 | 213 |
| SP71-6163 VA vs. IACSP95-5000 VA | 21 | 156 | 177 |

NA: no aphids (control); AA: aviruliferous aphids; VA: viruliferous aphids.

The infestation by aviruliferous and viruliferous aphids led mostly to the upregulation of genes in the SCYLV-susceptible cultivar SP71-6163. Sixty-three genes were upregulated and 7 were downregulated in response to aviruliferous aphids, while 23 were upregulated and 9 were downregulated in response to viruliferous aphids. In the tolerant cultivar IACSP95-5000, infestation by aviruliferous aphids led mainly to gene upregulation, with 589 upregulated and 64 downregulated genes. However, infection by viruliferous aphids led to the downregulation of several genes, with 27 upregulated and 138 downregulated genes. These results were also clearly observed in the comparisons between the two cultivars, in which a low number of DEGs (26) were found in the control condition. Additionally, in both cultivars, the number of DEGs in the comparison between controls and plants with aviruliferous aphids (70 in SP71-6163 and 653 in IACSP95-5000) was much greater than that in the comparisons between controls and plants with viruliferous aphids (32 in SP71-6163 and 165 in IACSP95-5000).

A more detailed investigation of DEG annotations and the results of Gene Ontology (GO) enrichment analyses further demonstrated the differences in the responses of each cultivar to both *M. sorghi* and SCYLV. REVIGO TreeMaps showing the biological processes predominating in each DEG set are shown in Figures 3, S3 and S4. In the response of SP71-6163 to aviruliferous aphids (Figure S3), terms associated with ‘ent-kaurene oxidation to kaurenoic acid’ and ‘negative regulation of transcription by RNA polymerase II’ were enriched. IACSP95-5000 seemed to activate more defense-specific pathways in response to *M. sorghi*. In this cultivar, two processes with a clear association with plant resistance to aphids were highly represented: the metabolism of flavonoids and the regulation of Rac protein signal transduction (Figure S4). The representative genes for these processes are Cluster_17139, encoding a C-type lectin domain containing 16A (CLEC16A) protein, and Cluster_45858, encoding afadin- and alpha-actinin-binding protein B, both of which were upregulated in response to *M. sorghi*. When the two cultivars were compared, processes associated with ‘innate immune response’, ‘regulation of histone H3-K9 methylation’ and ‘signal peptide processing’ were found to be enriched (Figure 3). The genes Cluster_10338 and Cluster_5506, annotated as a phytosulfokine receptor 2 and an E3 ubiquitin-protein ligase BRE1-like 2, respectively, were representative of the ‘innate immune response’ process and were upregulated in IACSP95-5000 infested with aviruliferous aphids.

**Figure 3.**
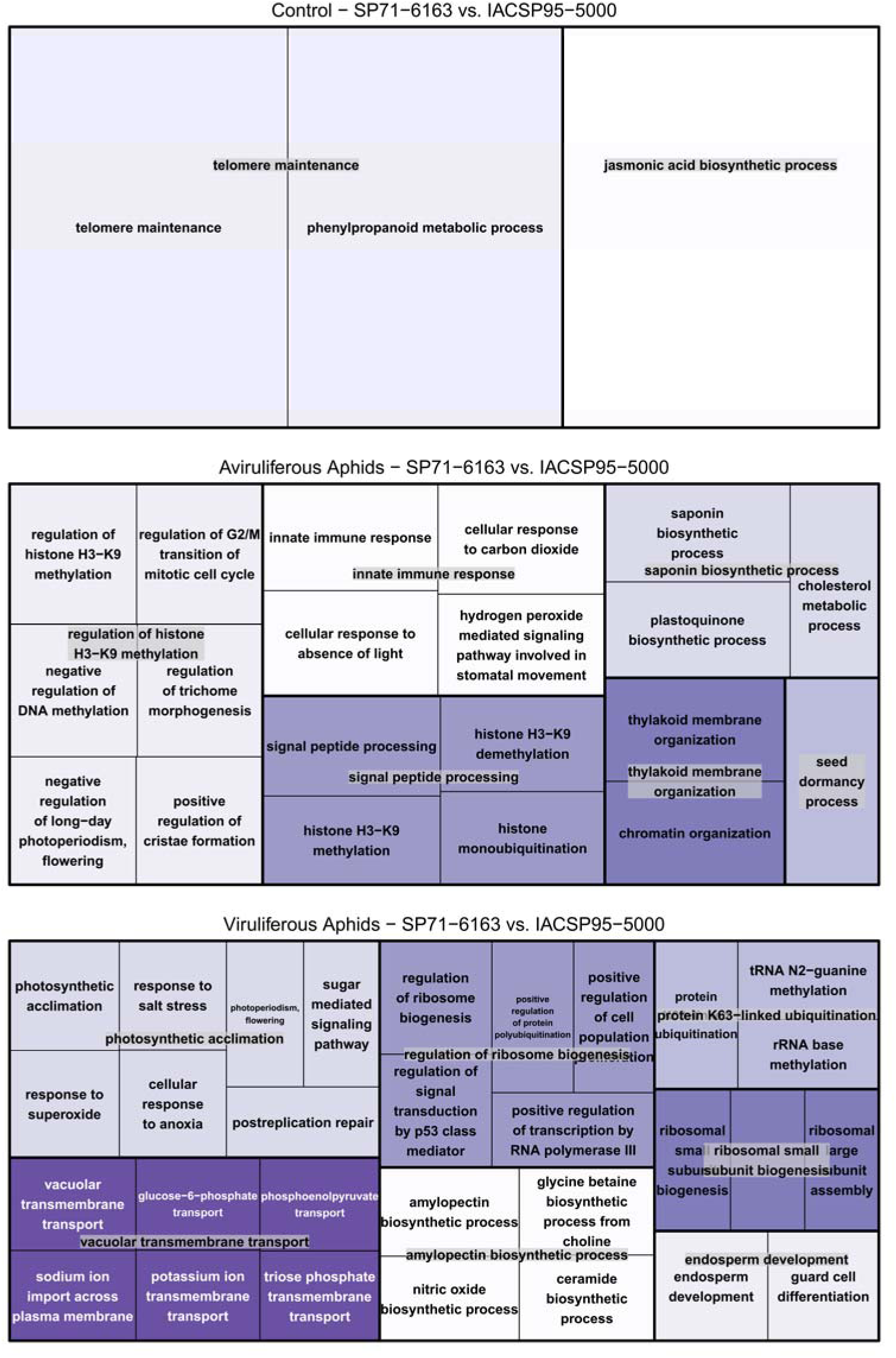
REVIGO representation of Gene Ontology categories enriched in the sets of differentially expressed genes from comparisons between SP71-6163 and IACSP95-5000 under the same treatments.

Differences were also clear when analyzing the response of the two cultivars to viruliferous aphids. In SP71-6163, ‘methionine biosynthetic process’ and ‘methylation’ categories were enriched (Figure S3). In IACSP95-5000, processes associated with the biogenesis of ribosomes constituted a high proportion of enriched GO terms, along with others such as the biosynthesis of amylopectin (Figure S4). Ribosome biogenesis was apparently altered in IACSP95-5000 in response to SCYLV, with the downregulation of a 25S rRNA (cytosine-C(5))-methyltransferase gene (Cluster 21838), and amylopectin synthesis was also affected, with differential expression of a soluble starch synthase (Cluster_1012). These two processes were also clearly represented when the two cultivars were compared, together with ‘photosynthetic acclimation’ and ‘vacuolar transmembrane transport’ (Figure 3).

In the comparison of IACSP95-5000 infested with aviruliferous and viruliferous aphids, several processes that were present in other comparisons also stood out. GO terms associated with ‘flavonoid transport from endoplasmic reticulum to plant-type vacuole’, for instance, were present in the comparisons between plants with viruliferous aphids and both the control and plants with aviruliferous aphids, with several DEGs identified in within-genotype comparisons also present in these comparisons. However, genes linked to these terms, such as Cluster_17139, were upregulated in response to *M. sorghi* alone and downregulated in response to SCYLV-viruliferous aphids. Notably, genes associated with the regulation of Rac protein signal transduction were upregulated in response to aviruliferous aphids but were not differentially expressed between the control plants and plants infested with viruliferous aphids. Similarly, GO terms associated with the biogenesis of ribosomes and amylopectin biosynthesis were identified via comparisons of plants exposed to viruliferous aphids with both the non-inoculated controls and plants exposed to aviruliferous aphids.

### 2.5. Gene coexpression network analyses

After additional filtering steps, transcript per million (TPM)-normalized data from 85,685 genes were used to construct a gene coexpression network via WGCNA. The pairwise Pearson correlation coefficients were increased to a soft threshold with β = 7, corresponding to an R^2^ of 0.798 and a mean connectivity of 536 (Figure S5). Using the unweighted pair group method with arithmetic mean (UPGMA) approach, 47 functional modules were defined in this network, with sizes ranging from 125 to 14,816 genes and an average size of 2,112 genes (Table S8 and Figure S6). Additionally, the genes with the highest connectivity in each module were identified as hubs. Three genes identified as hubs in their modules were also identified as DEGs. These genes were Cluster_21034 in the darkorange module, annotated as an L-type lectin domain-containing receptor kinase; Cluster_140592 in the ivory module, annotated as histone H4; and Cluster_39399 in the violet module, annotated as autophagy-related protein 16-1.

Some DEG sets were overrepresented in eight network modules, with some highly significant associations (Table S9). Notably, the three modules whose hub genes were also differentially expressed—darkorange, ivory, and violet—were among them. The most significant associations involved the modules darkorange, plum1 and violet, associated with comparisons including IACSP95-5000 infested with aviruliferous aphids, and magenta, associated with comparisons that included IACSP95-5000 infested with viruliferous aphids. These modules were enriched for various biological processes, as depicted in Figure S7. In the darkorange module, ‘regulation of DNA stability’ and ‘lipoxygenase pathway’ were the most prevalent categories. In the plum1 module, ‘regulation of glucose metabolic process’ and ‘flavonoid transport from endoplasmic reticulum to plant-type vacuole’ were predominant. In the violet module, ‘regulation of muscle cell differentiation’ and ‘protein urmylation’ were the most prevalent categories. In the magenta module, ‘shade avoidance’ and ‘copper ion transport’ prevailed. Other interesting processes enriched in other modules were ‘regulation of peroxidase activity’ and ‘regulation of ncRNA transcription’ in the ivory module and ‘regulation by virus of viral protein levels in host cell’ in the midnightblue module.

## 3. Discussion

Herein, *M. sorghi* was shown to be a vector of SCYLV in sugarcane. Until recently, this aphid species was considered restricted to Africa, but this view changed after the invasion of sorghum by the multilocus lineage (MLL)-F in the United States in 2013 (Bowling et al. 2016). Initially identified as *M. sacchari* (Nibouche et al. 2018), MLL-F was later identified by Nibouche et al. (2021) as a lineage of *M. sorghi.* These authors also presented evidence that despite having a preference for sorghum, this species also infests sugarcane. Many previous studies that focus on *Melanaphis* spp. do not mention species identification methodologies and often reference *M. sacchari*, but others have supported the hypothesis that *M. sorghi* is responsible for recent aphid outbreaks in sorghum in the Americas (Harris-Shultz et al. 2022; Balbi et al. 2023). Given the threat this aphid has posed in recent years, the evidence that it is capable of transmitting SCYLV is worrisome, especially given the potential detrimental effects of SCYLV on the host response to *M. sorghi*.

The DGE analysis performed here indicated that SCYLV-susceptible SP71-6163 has an incipient response to both SCYLV and *M. sorghi*, with very few genes activated or repressed in response to this virus and its vector. Its response to aviruliferous aphids led to the upregulation of Cluster_3551, encoding a leucine repeat-rich (LRR) receptor-like kinase that triggers plant innate immunity (Wang et al. 2021). However, the functions of most DEGs identified in comparisons involving only SP71-6163 were overall not consistent with what is commonly described for plant transcriptomic responses to sucking insects, which usually include secondary metabolic processes, jasmonic and salicylic acid biosynthesis, and signal transduction (Zogli et al. 2020), neither to viruses, such as RNA silencing and other pathways related to plant host defense and stress responses (Zanardo et al. 2019).

However, some of these processes were clearly and differentially represented in the responses of SCYLV-tolerant IACSP95-5000. In addition to the upregulation of the LRR kinase-encoding gene Cluster_3551, plants of this cultivar exposed to aviruliferous *M. sorghi* exhibited upregulation of genes associated with the biosynthesis of flavonoids, a class of chemically diverse secondary metabolites that are highly relevant in the defense of plants against insects (Treutter 2006). Genes involved in flavonoid biosynthesis have been previously shown to be upregulated in sorghum infested with *M. sacchari* (Kiani and Szczepaniec 2018; Tetreault et al. 2019). Here, genes encoding a CLEC16A protein (Cluster_17139), which has domains associated with intracellular membrane trafficking and the accumulation of flavonoids in plants (Ichino et al. 2014), and a flavonoid 3’-monooxygenase (Cluster_78445), which is involved in the biosynthesis of these compounds, were upregulated in response to *M. sorghi* in IACSP95-5000. Similarly, the signaling of Rac proteins, a class of Rho GTPases, may be enhanced in IACSP95-5000 through the upregulation of afadin- and alpha-actinin-binding proteins observed in plants exposed to *M. sorghi*. These proteins are also linked to the production of reactive oxygen species and thus to plant responses to biotic stresses (Torres 2010). Indeed, a Rac protein has been demonstrated to improve the defense responses of cotton plants against the aphid *Myzus persicae* (Yang et al. 2018).

IACSP95-5000 also responded very differently to infection by SCYLV. Notably, infection with this virus seemed to lead to the regulation of genes linked to the biogenesis of ribosomes and the biosynthesis of amylopectin. A representative upregulated gene associated with the biogenesis of ribosomes is an rRNA methyltransferase, which participates in rRNA methylation, a process that optimizes ribosomal function (Zou et al. 2020). In general, RNA viruses are known to induce heterogeneous ribosome production (Wang et al. 2022), and differential expression of ribosomal protein genes has been reported in studies analyzing plant responses to SCYLV and other plant viruses (Shabbir et al. 2022; Zhang et al. 2022).

The synthesis of amylopectin may also be affected by the modulation of the expression of starch synthase, an enzyme that plays a key role in the synthesis of this polysaccharide (Irshad et al. 2021) and has even stronger links with infection by SCYLV. Amylopectin is the major component of starch, composing ca. 80% of this polymeric carbohydrate in sugarcane (Figueira et al. 2011; Alves et al. 2014). The most notorious effects of infection by SCYLV in sugarcane relate to the metabolism and transport of carbohydrates, one of which is the increased content of starch in the leaves, especially in tolerant or resistant varieties (Lehrer et al. 2007, 2010; Yan et al. 2008). Thus, the differential regulation of genes affecting the synthesis of the main constituents of starch in response to SCYLV could be associated with its accumulation during infection.

One of the most notable results observed here was the fewer DEGs identified in both sugarcane cultivars exposed to viruliferous aphids than in those exposed to aviruliferous aphids. This is an indication that infection by SCYLV has a negative effect on the plant’s response to its vector, an occurrence that has been observed in other members of the *Polerovirus* genus, including potato leafroll virus (PLRV) and turnip yellows virus (TuYV) (Patton et al. 2020; Krieger et al. 2023). Krieger et al. (2023) recently demonstrated that TuYV has a suppressive effect on the defense mechanisms of *Arabidopsis thaliana* against *M. persicae* to increase its transmission efficiency. It is possible that a similar phenomenon occurs in the sugarcane-*M. sorghi*-SCYLV system, in which the virus seemingly suppresses the activation of pathways leading to the biosynthesis of compounds that would otherwise be upregulated in response to the aphid alone.

Additionally, a gene coexpression network revealed broader biological processes associated with sugarcane responses to *M. sorghi* and SCYLV. The darkorange module had an overrepresentation of DEGs from comparisons involving IACSP-95-5000 infested with aviruliferous aphids, among which was the top hub of this module. This gene encodes an L-type lectin domain-containing receptor kinase, a class of protein that has been shown to act as an immune receptor in plants and activate downstream defense pathways (Wang and Bouwmeester 2017). The recognition of attack by *M. sorghi* by such a kinase could lead to the activation of defense-related processes such as the lipoxygenase pathway, which was enriched in the darkorange module. Lipoxygenases are a family of enzymes with diverse biological roles, including the response to biotic and abiotic stresses. Some lipoxygenases are differentially expressed in sorghum in response to *M. sacchari* feeding (Shrestha et al. 2021), so their involvement in the response of sugarcane to *M. sorghi* is biologically plausible.

A similar scenario was observed for the violet module, which also had an overrepresentation of DEGs associated with IACSP-95-5000 infested with aviruliferous aphids, including the top hub of the module. This gene encodes an autophagy-related protein, a class of proteins that modulates cell death in plants under biotic stresses, including aphid feeding (Hao et al. 2023). Consistent with this finding, the violet module was enriched with several biological processes linked with autophagy, such as cell killing and protein catabolism, which may be enhanced in IACSP95-5000 in response to *M. sorghi*. Crucially, the hubs of the violet and darkorange modules were not differentially expressed in response to viruliferous aphids, reinforcing the hypothesis that SCYLV disrupted the associated immune responses.

The plum1 module, which also showed high overrepresentation of DEGs identified in comparisons involving IACSP95-5000 infested with aviruliferous aphids, is also worthy of further discussion. In addition to the terms associated with flavonoid transport, whose role in response to aphids has been discussed, this module was enriched with terms linked with the metabolism of glucose. In sorghum, *M. sacchari* feeding is known to affect the content of saccharides and increase the content of glucose, which has also been correlated with increased damage caused by this aphid (Knoll et al. 2021; Cardona et al. 2023). Finally, the midnightblue module, which was associated with comparisons involving IACSP95-5000 infested with viruliferous aphids, was enriched with ‘regulation by virus of viral protein levels in host cell’ terms, which can be linked to the plant response to SCYLV. These results revealed further potential effects of *M. sorghi* feeding and infection with SCYLV that were not captured in the DGE analysis alone, demonstrating the ability of gene coexpression network methodologies to expand the scope of transcriptomic analyses.

The potential of SCYLV to affect host response to *M. sorghi* highlights the need for further investigations on the incidence of SCYLV in both this aphid and in the crops it infests, which could exacerbate the direct damage the aphid inflicts. Research on the percentage of *Melanaphis* spp. aphids carrying SCYLV has never been carried out on any crop. Studies analyzing aphids vectoring other poleroviruses have revealed incidences of up to 100% (Singh et al. 1997; Thango 2019). SCYLV is often highly prevalent in commercial sugarcane fields (Lehrer et al. 2008; Amata et al. 2016; Filloux et al. 2018) and could serve as an inoculum source for sorghum fields, where the incidence of this virus has thus far been shown to be relatively low (Boukari et al. 2021; Viswanathan et al. 2023). Recent research has shown that *M. sacchari* feeding on different sugarcane cultivars infected with SCYLV present differences in their behavior and biology (Bertasello et al. 2021). Thus, it would also be valuable to evaluate the performance and behavior of *Melanaphis* spp. on SCYLV-infected and virus-free plants to assess how the biology of these aphids is affected by the virus, which is consistent with the findings reported herein.

## 4. Conclusions

This study revealed that *M. sorghi* is capable of vectoring SCYLV, posing an additional threat to the crops currently impacted by this aphid. Moreover, these findings indicate that infection by SCYLV has a negative effect on plant defense responses to its vector, leading to shifts in the expression of genes involved in pathways modulated in response to feeding by *M. sorghi*. Contrasting responses were observed in a comparison between SCYLV-susceptible and SCYLV-tolerant sugarcane genotypes, indicating that susceptibility may result from a combination of ineffective responses to both SCYLV and *M. sorghi*. This highlights the need for further investigations on the impacts of SCYLV on its vectors and hosts and corresponding implications for virus epidemiology.

## 5. Methods

### 5.1. Plant material and virus isolate

Two Brazilian hybrid sugarcane cultivars were used in the experiment. SP71-6163 is efficiently infected by SCYLV, develops severe symptoms of YLD and is strongly affected by the virus in terms of yield (Vega et al. 1997; Rodrigues 2015); therefore, it was chosen as an SCYLV-susceptible genotype. Despite being a highly suitable host to *M. sacchari* and exhibiting SCYLV infection rates similar to those of SP71-6163 (Rodrigues 2015; Bertasello et al. 2021), IACSP95-5000 has not been reported to exhibit YLD symptoms or high yield losses (M.G.A. Landell, personal communication); for these reasons, this cultivar was selected as an SCYLV-tolerant genotype. Virus-free plants of the two cultivars were generated by meristem micropropagation at the Instituto Agronômico de Campinas Advanced Centre for Sugarcane Research and Development’s biofactory.

Plants clonally propagated from an SCYLV-infected SP71-6163 plant from Ribeirão Preto, Brazil (Pimenta et al. 2021), were used as a source of inoculum. Total RNA was extracted from the leaf of one plant following a protocol adapted from Kistner and Matamoros (2005). RNA was quantified on a NanoDrop 2000 spectrophotometer and subjected to electrophoresis on a 1% denaturing agarose gel stained with ethidium bromide for an integrity check. RT was performed with the ImProm-II Reverse Transcription System Kit (Promega, Madison, USA). Phusion High-Fidelity PCR Master Mix (New England BioLabs, Ipswich, USA) was used to amplify overlapping fragments covering the complete SCYLV genome. Primers were designed using Primer3Plus v. 3.3.0 (Untergasser et al. 2007); details are provided in Table S9. PCR products were purified with polyethylene glycol and sequenced on a 3500xL Genetic Analyzer (Applied Biosystems, Foster City, USA).

Geneious v. 7.1.3 software (Kearse et al. 2012) was used for Sanger sequencing data handling. Low-quality bases and primer sequences were manually trimmed, and reads were used to produce a de novo assembly of a contiguous complete genomic sequence. The complete genomes of 101 SCYLV isolates from 17 different countries, as well as the PLRV reference genome (Table S10), were retrieved from GenBank, pairwise-aligned using ClustalW v. 2.1 (Thompson et al. 2003) and trimmed to the length of the shortest sequences. Pairwise Hamming distances (Hamming 1950) were calculated on UGENE v. 48.1 (Okonechnikov et al. 2012). The presence of recombinants was determined using the Recombination Detection Program v. 5.33 (RDP5) (Martin et al. 2021) using seven different methods with default parameters: RDP, GENECONV, BOOTSCAN, MAXCHI, CHIMAERA, SISCAN and 3SEQ. Only recombination events detected by at least five methods and showing significant support with a Bonferroni-corrected *p* value cutoff of 0.05 were considered valid (Pagán and Holmes 2010). Recombinant isolates were removed for subsequent analyses, and MEGA11 v. 11.0.11 (Tamura et al. 2021) was used to find the best-fit DNA/protein model based on the Bayesian information criterion (BIC) score. The model with the lowest BIC score was used to construct a maximum likelihood tree rooted by the *Polerovirus* type species *Potato leafroll virus* (PLRV) genome via the same software.

Bootstrap procedures with 1,000 replicates were used to generate standard error estimates, and the resulting tree was visualized with FigTree v. 1.4.4 218 software (Rambault 2018).

### 5.2. Aphid rearing and identification

*Melanaphis* sp. aphids were obtained from a sugarcane field in Ribeirão Preto, Brazil, and reared on detached sugarcane leaf fragments in test tubes with 1% agar kept in a plant growth chamber at 30°C under 12-hour light/dark cycles. A colony of aviruliferous aphids was obtained by transferring individual adult aphids onto virus-free sugarcane leaves and transferring newborn nymphs to separate virus-free leaves. To ensure the effectiveness of the process, nymph transfer was repeated for several aphid generations, and the colonies obtained were tested for the presence of SCYLV by RT[PCR. For this purpose, RNA was extracted from pools of five aphids using TRIzol (Invitrogen, Carlsbad, USA), quantified on a NanoDrop spectrophotometer and converted into cDNA using the ImProm-II Reverse Transcription System Kit. cDNA was used for PCR diagnosis as described by Gonçalves et al. (2002). Prior to inoculation experiments, a viruliferous colony was obtained by transferring aphids to SCYLV-infected SP71-6163 plants kept in aphid-proof cages. Aphids were allowed an acquisition access period on infected plants of at least 48 hours for virus acquisition.

While *M. sacchari* is the most common *Melanaphis* species that infests sugarcane, *M. sorghi* has been recently reported in Brazil (Nibouche et al. 2021; Harris-Shultz et al. 2022). Because the two species are very similar morphologically, a molecular diagnostic marker was used to determine the species of the established colony. DNA was extracted from a pool of five aphids using a protocol adapted from Sunnucks and Hales (1996). The primers MelF2 (GTGCTTATTGTCGCTGCTGG) and MelR2 (TTGTCTCCGGGAACAGCTTC) were designed to amplify a 683-bp fragment of the *EF1-*α gene harboring five polymorphic sites in *M. sacchari* and *M. sorghi*, one of which is sufficient to distinguish the two species (Nibouche et al. 2021). PCR was performed using the Phusion High-Fidelity PCR Master Mix and an annealing temperature of 60°C. The PCR products were purified using polyethylene glycol and sequenced on a 3500xL Genetic Analyzer. Geneious was used to manually trim low-quality bases and primer sequences, perform pairwise alignment of forward and reverse reads, and analyze the nucleotides present at the polymorphic sites.

### 5.3. Inoculation experiment and sample processing

The inoculation experiment was performed in a plant growth chamber kept at 30°C under 12-hour light/dark cycles; three biological replicates consisting of 11-month-old plants were used for each experimental condition. Clip cages were positioned on the median region of the top visible dewlap leaf of each plant, in which five viruliferous or aviruliferous adult aphids were placed; for aphid-free control plants, no aphids were used. After inoculation access periods (IAPs) of 4, 24 or 48 hours, aphids were removed, and whole leaves were collected, flash-frozen in liquid nitrogen and stored at −80°C until processing.

Total RNA was extracted from the ~10 cm leaf section around the clip cage placement site, quality was checked, and RT was performed as previously described. cDNA was used to confirm infection and estimate the SCYLV titer by a qPCR-based relative quantification method detailed by Pimenta et al. (2021). The normality and variance of the viral quantification data were verified by Shapiro[Wilk and F tests, respectively, and differences in quantification between genotypes were assessed by an unpaired t test in R software (R Core Team 2011).

RNA from selected samples was used to prepare RNA-Seq libraries using the TruSeq Stranded mRNA Library Prep Kit (Illumina, San Diego, USA). After their fragment size profiles were verified on an Agilent 2100 Bioanalyzer (Agilent Technologies, Palo Alto, USA), the libraries were quantified on a Qubit 3.0 fluorometer (Thermo Scientific, Wilmington, USA), pooled and requantified. A 2×150 bp paired-end sequencing library was prepared with the NovaSeq 6000 SP Reagent Kit v1.5 (Illumina, San Diego, USA) and sequenced in two lanes of a NovaSeq 6000 instrument (Illumina, San Diego, USA).

### 5.4. Reference transcriptome assembly and annotation

In addition to the data generated for the samples from the present study, previously available RNA-Seq data were included in the analyses to contribute to the assembly of a representative transcriptomic reference and to the construction of a robust gene coexpression network.

These consisted of data from leaf samples collected from twelve diverse *Saccharum* genotypes generated by Correr et al. (2020). The quality of all RNA-Seq data was assessed using FastQC (Andrews 2010). Trimmomatic v.0.39 (Bolger et al. 2014) was used for read trimming, using a 5-bp sliding window with a minimum average Phred quality score of 20 and removing reads shorter than 75 bp.

Filtered reads were aligned to the genomes of *S. officinarum* (GenBank GCA_020631735.1) and *S. spontaneum* (GenBank GCA_022457205.1) using STAR v. 2.7.3 (Dobin et al. 2013). These genomes are allele-resolved, and each haplotype was treated as a different reference. Thus, twelve different alignments were performed for each sample, representing the eight haplotypes of *S. officinarum* and the four of *S. spontaneum*. After comparative mapping, the resulting files were sorted based on their genomic positions using SAMtools v. 1.12 (Li et al. 2009) and used to define twelve distinct transcriptomes via Stringtie v. 2.1.6 (Pertea et al. 2015). CD-HIT v. (Fu et al. 2012) was then used to reduce redundancies in the reference. This software was first used for the transcriptomes of each species to combine genes from individual haplotypes and subsequently to combine the transcriptomes generated for the two species. This final reference was evaluated with BUSCO v. 5.0.0 (Simão et al. 2015) using datasets of conserved orthologs from Eukaryota and Viridiplantae. It was then annotated with Trinotate v. 4.0.1 (Bryant et al. 2017) by performing a homology search of sequences on UniProt release 2024_1 (UniProt Consortium 2019).

### 5.5. Differential gene expression analyses

Salmon v.1.1.0 software (Patro et al. 2017) was used for transcript quantification via selective alignment of the sequencing reads and correction for fragment-level GC biases. The tximport R package v. 1.26.1 (Soneson et al. 2015) was used to import quantification data into R software. The edgeR package v 3.40.2 (Robinson et al. 2010) was used to normalize quantifications by CPM. For the differential gene expression analysis, in which only samples generated in the present work were used, specific filters were used. First, to remove genes with low expression across treatments, only genes with a sum of CPMs ≥ 3 in all samples of a treatment were retained.

Next, genes with inconsistent expression across replicates of each treatment were filtered out. To achieve this goal, paired t tests were performed to identify genes with significant (*p* < 0.05) differences in CPM values across the six lanes encompassing each treatment, which were discarded. Subsequently, the gene sets for each biological condition were combined according to the nine comparisons performed: (1) SP71-6163 Control vs. SP71-6163 Aviruliferous Aphids; (2) SP71-6163 Control vs. SP71-6163 Viruliferous Aphids; (3) SP71-6163 Aviruliferous Aphids vs. SP71-6163 Viruliferous Aphids; (4) IACSP95-5000 Control vs. IACSP95-5000 Aviruliferous Aphids; (5) IACSP95-5000 Control vs. IACSP95-5000 Viruliferous Aphids; (6) IACSP95-5000 Aviruliferous Aphids vs. IACSP95-5000 Viruliferous Aphids; (7) SP71-6163 Control vs. IACSP95-5000 Control; (8) SP71-6163 Aviruliferous Aphids vs. IACSP95-5000 Aviruliferous Aphids; and (9) SP71-6163 Viruliferous Aphids vs. IACSP95-5000 Viruliferous Aphids.

DGE analyses were performed with the R package DESeq2 v. 1.38.3 (Love et al. 2014) using a false discovery rate (FDR)-corrected *p* value of 0.05 and a log_2_(FC) value of 1 as thresholds for the delimitation of DEGs. The UpSetR R package (Conway et al. 2017) was used to investigate overlaps between sets of DEGs and generate an UpSet plot. GO enrichment analysis was performed on each DEG set with the topGO R package v. 2.50.0 (Alexa et al. 2006) using Fisher’s test with α = 0.05. REVIGO v. 1.8.1 (Supek et al. 2011) was used for the visualization and analysis of the enriched GO categories.

### 5.6. Gene coexpression network

A gene coexpression network was constructed using data generated in the present study as well as those from Correr et al. (2020). Quantification data were filtered to retain genes with a CPM ≥ 1 in all samples from at least one treatment. The edgeR R package was used to normalize quantifications by TPM, and the WGCNA R package v. 1.72.5 (Langfelder and Horvath 2008) was used to construct the gene coexpression network. Pairwise Pearson correlations of TPM values considering a power function fitting scale-free independence were used. A soft threshold value corresponding to an R^2^ value of approximately 0.8 was used to approximate a scale-free topology model. Functional modules in the network were defined using UPGMA based on a topological overlap matrix and dynamic dendrogram pruning based only on the dendrogram. Hubs, defined as the genes with the highest connectivity in each module, were identified with the dedicated WGCNA function.

Fisher’s exact test with an FDR correction with α = 0.05, performed in R, was used to verify whether DEG sets identified in the DGE analyses were overrepresented in any of the modules of the network. GO enrichment analysis was performed on the genes included in these modules with topGO using Fisher’s test with α = 0.05, and enriched GO terms were visualized with REVIGO.

### 5.7. RT***__***qPCR validation

RNA-Seq libraries were validated by RT[qPCR. Primers to amplify eight DEGs and one gene stably and highly expressed across all samples were designed using Primer3Plus with parameters optimized for qPCR. Additionally, primers designed to amplify one common sugarcane control gene, *GAPDH* (Iskandar et al. 2004), were synthesized. The sequences of all the qPCR primers can be found in Table S11. cDNA was synthesized using the QuantiTect Reverse Transcription Kit (QIAGEN, Hilden, Germany) following the manufacturer’s instructions. qPCRs were performed using the iTaq Universal SYBR Green Supermix (Bio-Rad, Philadelphia, USA) on a Bio-Rad CFX384 Touch detection system (Bio-Rad, Philadelphia, USA) according to the manufacturer’s cycling recommendations. The mean Cq values of the two endogenous controls were used to correct the DEG Cq values. All reactions were performed using three technical replicates. Linear models correlating the CPM and ΔCq values were fitted and plotted in R using the ggplot2 (Wickham 2016) and ggpubr (Kassambara 2023) packages.

## Declarations

### Ethics approval and consent to participate

Not applicable.

### Consent for publication

Not applicable.

### Availability of data and materials

The RNA sequencing data generated by the present work have been deposited in SRA under the BioProject ID PRJNA1125051. The complete genome of the BRA-SP1 isolate and the *M. sorghi EF1-*α partial sequence have been deposited in GenBank under accession numbers PP796534 and PP796535, respectively. Table S6 is available on FigShare under the DOI 10.6084/m9.figshare.27236814.

## Funding

This work was supported by grants from the São Paulo Research Foundation (FAPESP), the Conselho Nacional de Desenvolvimento Científico e Tecnológico (CNPq, grant 424050/2016-1), and the Coordenação de Aperfeiçoamento de Pessoal de Nível Superior (CAPES, Computational Biology Program). RJGP received MSc and PhD fellowships from FAPESP (grants 2018/18588-8 and 2019/21682-9). AHA received a PhD fellowship from FAPESP (grant 2019/03232-6).

### Authors’ contributions

MCG, LRP and APS conceived the project and designed the experiments. MNGM generated the virus-free plants. RJGP and DAS performed further experimental procedures. RJGP and AHA analyzed the data. RJGP interpreted the results and wrote the manuscript. All the authors have read and approved the manuscript.

## List of abbreviations

IAP: Inoculation access period
BIC: Bayesian information criterion
CLEC16A: C-type lectin domain containing 16A
CPM: Counts per million
DGE: Differential gene expression
DEG: Differentially expressed gene
EF1-α: Elongation factor 1-alpha
LRR: Leucine repeat-rich
GAPDH: Glyceraldehyde-3-phosphate dehydrogenase
GO: Gene Ontology
MLL: Multilocus lineage
PLRV: Potato leafroll virus
RT qPCR: Reverse transcription followed by quantitative polymerase chain reaction
SCYLV: Sugarcane yellow leaf virus
TPM: Transcripts per million
TuYV: Turnip yellows virus
UPGMA: Unweighted pair group method with arithmetic mean
YLD: Yellow leaf disease

## Acknowledgements

We thank Silvana Creste for the use of the IAC biofactory for the production of virus-free sugarcane plants, Maicon Volpin for assistance in aphid rearing and greenhouse work and Aline C. L. Moraes for assistance in constructing and sequencing the RNA-Seq library.

