## Supplementary Material for "Sugarcane yellow leaf virus impairs the transcriptomic defense responses of sugarcane to its new aphid vector *Melanaphis sorghi*"

### Supplementary Tables

#### Table S1: Recombination events detected between sugarcane yellow leaf virus isolates.

| Event Number | Recombinant | Minor Parent | Major Parent | Method *p* value | | | | | | |
|---|---|---|---|---|---|---|---|---|---|---|
| | | | | RDP | GENECONV | Bootscan | Maxchi | Chimaera | SiSscan | 3Seq |
| 1 | GT-01a | GT-02b | GT-01b | 2.03E-42 | 2.94E-38 | 8.68E-42 | 2.27E-22 | 2.54E-22 | 2.84E-28 | 9.38E-41 |
| 2 | PI_157033 | chn1 | BR-03b | 6.01E-38 | 1.98E-36 | 1.57E-35 | 1.85E-08 | 2.04E-08 | 1.83E-13 | 8.50E-19 |
| 3 | Sorg1_1 | Sorg2_2 | IND3 | 9.93E-23 | 3.62E-26 | 5.15E-24 | 1.70E-19 | 3.27E-15 | 1.68E-24 | 3.50E-18 |
| 4 | Sorg3_3 | CO-06 | GT-07 | 7.06E-21 | 7.84E-23 | 5.22E-23 | 6.68E-12 | 9.39E-11 | 8.60E-16 | 1.71E-10 |
| 5 | MU-AB193 | BR-06-Leaf | REU-YL2 | 1.13E-17 | 6.65E-16 | 2.57E-13 | 1.49E-13 | 2.81E-14 | 1.91E-16 | 2.50E-22 |
| 6 | GT-01a | GT-02a | BRA-YL1 | 2.87E-19 | 3.30E-17 | 2.19E-02 | 2.18E-09 | 6.73E-09 | 1.14E-12 | 1.22E-11 |
| 7 | GT-02b | COL-YL1a | chn1 | 4.12E-03 | 3.88E-02 | NS | 9.98E-08 | 4.84E-08 | 1.39E-26 | 2.36E-18 |
| 8 | MU-AB193 | BR-02 | REU-YL2 | 6.94E-19 | 2.94E-17 | NS | 1.80E-07 | 1.62E-07 | 5.54E-09 | 2.13E-14 |
| 9 | MU-SC1233 | US-FL06 | MU-AB193 | 1.11E-12 | 1.24E-11 | 7.00E-08 | 5.85E-09 | 1.27E-08 | 1.19E-09 | 3.71E-18 |
| 10 | Sorg1_1 | GT-01b | GT-02a | 2.58E-17 | 5.28E-16 | NS | 1.34E-06 | 3.79E-06 | 6.05E-31 | 2.44E-11 |
| 11 | Sorg2_2 | CO-08 | GT-02a | 2.48E-13 | 5.28E-07 | NS | 4.11E-05 | 1.61E-03 | 1.07E-06 | 2.43E-73 |
| 12 | Sorg3_3 | MU-SC1233 | GT-03 | 3.23E-13 | 1.22E-10 | NS | 1.67E-05 | 9.74E-08 | NS | 4.40E-07 |
| 13 | GT-02a | CUB-YL1a | chn1 | 3.83E-08 | NS | NS | 2.20E-07 | 1.79E-07 | 2.83E-04 | 2.44E-11 |
| 14 | GT-01a | GT-07 | GT-01b | 5.38E-11 | 1.34E-09 | NS | 1.78E-04 | 1.89E-04 | 3.06E-07 | 6.93E-08 |
| 15 | MU-SC1233 | GT-01a | MU-AB110 | 1.02E-09 | 5.16E-08 | NS | 1.51E-04 | 1.82E-04 | 2.43E-14 | 3.12E-09 |
| 16 | MU-SC1233 | US-FL03 | MU01-b | 4.67E-11 | 1.83E-09 | 3.79E-03 | 1.33E-05 | 1.23E-05 | 8.86E-08 | 1.19E-09 |
| 17 | GT-01a | GT-02a | GT-01b | 4.10E-09 | 4.65E-06 | NS | 2.65E-02 | NS | 3.29E-11 | 1.33E-04 |
| 18 | Sorg3_3 | US-LA09 | GT-02a | 2.11E-06 | 9.75E-08 | 6.43E-09 | 1.04E-04 | NS | NS | 7.01E-04 |
| 19 | GT-01a | GT-02b | US-FL01b | 6.56E-09 | 1.08E-07 | NS | 1.03E-03 | 2.34E-02 | NS | 2.88E-05 |
| 20 | Sorg3_3 | IND | chn1 | 1.74E-05 | 6.15E-08 | NS | 1.43E-05 | 2.29E-05 | 8.05E-35 | 1.04E-04 |
| 21 | GT-02a | US-HI02 | Sorg3_3 | 1.21E-07 | 2.07E-07 | 8.97E-06 | 2.40E-03 | NS | 1.23E-03 | 3.96E-09 |
| 22 | GT-02a | GT-01b | Sorg3_3 | 1.90E-09 | 1.25E-07 | 7.05E-07 | 5.83E-03 | 2.53E-02 | 2.45E-04 | 6.27E-06 |
| 23 | IND3 | chn1 | CBLK97154 | 8.07E-06 | 1.75E-06 | 2.00E-07 | 3.07E-02 | NS | 2.06E-02 | 6.07E-04 |
| 24 | IND1 | IND4 | CO-02 | 5.24E-06 | 1.29E-05 | 2.18E-08 | 1.42E-04 | 9.11E-05 | 2.58E-08 | NS |
| 25 | Sorg2_2 | IND | IND3 | 7.81E-06 | 8.66E-03 | 3.88E-02 | 3.30E-03 | NS | 7.60E-33 | 2.40E-02 |
| 26 | MU01-a | chn1 | US-FL02 | 2.90E-08 | NS | 2.51E-04 | 5.15E-06 | 2.70E-04 | NS | 1.82E-75 |
| 27 | ZA-02 | PER-YL1a | CN-02 | 3.17E-05 | NS | 2.36E-02 | NS | 3.87E-02 | 2.25E-07 | 1.90E-02 |

NS: Nonsignificant.

**Table S2. Number and percentage of sugarcane plants infected by sugarcane yellow leaf virus at different inoculation access periods (IAPs) as detected by RT-qPCR.**

| Condition (IAP) | Plants infected | |
|---|---|---|
| | SP71-6163 | IACSP95-5000 |
| No aphids – Control | 0/3 (0%) | 0/3 (0%) |
| Aviruliferous aphids (4 hours) | 0/3 (0%) | 0/3 (0%) |
| Aviruliferous aphids (24 hours) | 0/3 (0%) | 0/3 (0%) |
| Aviruliferous aphids (48 hours) | 0/3 (0%) | 0/3 (0%) |
| Viruliferous aphids (4 hours) | 1/3 (33%) | 0/3 (0%) |
| Viruliferous aphids (24 hours) | 1/3 (33%) | 3/3 (100%) |
| Viruliferous aphids (48 hours) | 3/3 (100%) | 3/3 (100%) |

**Table S3: Number of raw and trimmed reads and alignment rate for each sample.**

| Sample | Raw reads | Trimmed reads | Alignment rate |
|---|---|---|---|
| SP71-6163 NA R1 | 47,170,812 | 39,202,794 | 89.11% |
| SP71-6163 NA R2 | 61,952,065 | 51,932,696 | 89.46% |
| SP71-6163 NA R3 | 50,054,898 | 42,787,473 | 89.13% |
| SP71-6163 AA 48h R1 | 38,530,843 | 32,647,033 | 89.37% |
| SP71-6163 AA 48h R2 | 54,333,763 | 45,292,467 | 89.53% |
| SP71-6163 AA 48h R3 | 56,113,062 | 47,929,930 | 89.49% |
| SP71-6163 VA 48h R1 | 122,450,434 | 103,727,067 | 89.6% |
| SP71-6163 VA 48h R2 | 89,158,811 | 71,569,626 | 89.54% |
| SP71-6163 VA 48h R3 | 50,910,761 | 41,540,905 | 88.9% |
| IACSP95-5000 NA R1 | 42,890,947 | 35,127,902 | 88.7% |
| IACSP95-5000 NA R2 | 44,537,543 | 35,996,995 | 90.58% |
| IACSP95-5000 NA R3 | 29,127,601 | 19,745,412 | 89.15% |
| IACSP95-5000 AA 48h R1 | 35,386,474 | 27,714,867 | 89.18% |
| IACSP95-5000 AA 48h R2 | 41,757,453 | 33,219,118 | 88.89% |
| IACSP95-5000 AA 48h R3 | 43,301,828 | 35,730,476 | 88.75% |
| IACSP95-5000 VA 48h R1 | 51,119,159 | 45,165,460 | 90.33% |
| IACSP95-5000 VA 48h R2 | 56,360,566 | 47,095,241 | 89.16% |
| IACSP95-5000 VA 48h R3 | 45,841,110 | 37,682,530 | 90.14% |

NA: no aphids (control), AA: aviruliferous aphids, VA: viruliferous aphids.

**Table S4: Number and percentage of conserved orthologs from Eukaryota and Viridiplantae found in the transcriptomic reference.**

| BUSCOs | Eukaryota | Viridiplantae |
|---|---|---|
| Complete and single-copy | 56 (22.0%) | 82 (19.3%) |
| Complete and duplicated | 197 (77.3%) | 336 (79.1%) |
| Fragmented | 1 (0.4%) | 5 (1.2%) |
| Missing | 1 (0.3%) | 2 (0.4%) |
| Total searched | 255 | 425 |

**Table S5. Number of genes excluded and retained in the filtering of gene quantification data, according to the comparison of the differential gene expression analysis.**

| Condition 1 | Condition 2 | Excluded Condition 1 | Excluded Condition 2 | Union 1+2 | Retained |
|---|---|---|---|---|---|
| SP71-6163 NA | SP71-6163 AA | 9,988 | 9,081 | 17,821 | 67,864 |
| SP71-6163 NA | SP71-6163 VA | 9,988 | 8,179 | 17,105 | 68,580 |
| SP71-6163 AA | SP71-6163 VA | 9,081 | 8,179 | 16,351 | 69,334 |
| IACSP95-5000 NA | IACSP95-5000 AA | 12,863 | 13,386 | 23,645 | 62,040 |
| IACSP95-5000 NA | IACSP95-5000 VA | 12,863 | 11,371 | 22,172 | 63,513 |
| IACSP95-5000 AA | IACSP95-5000 VA | 13,386 | 11,371 | 22,602 | 63,083 |
| SP71-6163 NA | IACSP95-5000 NA | 9,988 | 12,863 | 20,998 | 64,687 |
| SP71-6163 AA | IACSP95-5000 AA | 9,081 | 13,386 | 20,873 | 64,812 |
| SP71-6163 VA | IACSP95-5000 VA | 8,179 | 11,371 | 8,179 | 77,506 |

NA: no aphids (control), AA: aviruliferous aphids, VA: viruliferous aphids.

**Table S7. Size and hub gene identified for each module defined in the gene coexpression network.**

| Module | Size | Hub |
|---|---|---|
| black | 6,373 | Cluster_186271 |
| blue | 7,373 | Cluster_128389 |
| brown | 4,845 | Cluster_201021 |
| brown4 | 365 | Cluster_38346 |
| cyan | 1,407 | Cluster_57437 |
| darkgreen | 1,071 | Cluster_85082 |
| darkgrey | 1,852 | Cluster_81132 |
| darkmagenta | 611 | Cluster_101653 |
| darkolivegreen | 1,239 | Cluster_694 |
| darkorange | 788 | Cluster_21034 |
| darkorange2 | 125 | Cluster_83089 |
| darkred | 1,147 | Cluster_43164 |
| darkturquoise | 1,387 | Cluster_120266 |
| floralwhite | 426 | Cluster_25418 |
| green | 5,401 | Cluster_65119 |
| greenyellow | 1,568 | Cluster_142263 |
| grey60 | 1,023 | Cluster_65026 |
| ivory | 581 | Cluster_140592 |
| lightcyan | 1,022 | Cluster_67804 |
| lightcyan1 | 816 | Cluster_120861 |
| lightgreen | 829 | Cluster_126796 |
| lightsteelblue1 | 814 | Cluster_122869 |
| lightyellow | 1,073 | Cluster_2460 |
| magenta | 3,224 | Cluster_85708 |
| mediumpurple3 | 559 | Cluster_151866 |
| midnightblue | 1,637 | Cluster_194635 |
| orange | 1,992 | Cluster_193758 |
| orangered4 | 468 | Cluster_23321 |
| paleturquoise | 938 | Cluster_176389 |
| pink | 4,195 | Cluster_122592 |
| plum1 | 367 | Cluster_128525 |
| purple | 3,132 | Cluster_81914 |
| red | 3,753 | Cluster_114480 |
| royalblue | 992 | Cluster_9687 |
| saddlebrown | 1,304 | Cluster_43540 |
| salmon | 2,729 | Cluster_159121 |
| sienna3 | 828 | Cluster_172158 |
| skyblue | 826 | Cluster_138835 |
| skyblue3 | 818 | Cluster_54349 |
| steelblue | 626 | Cluster_158438 |
| tan | 1,898 | Cluster_149097 |
| turquoise | 14,816 | Cluster_192933 |
| violet | 567 | Cluster_39399 |
| white | 1,653 | Cluster_42571 |
| yellow | 8,121 | Cluster_26477 |
| yellowgreen | 731 | Cluster_17543 |

**Table S8. Sets of differentially expressed genes overrepresented in the coexpression network modules, along with module size, expected and observed counts and false discovery rate-adjusted *p* values.**

| Comparison | Module | Module size | Expected count | Observed count | *p* value |
|---|---|---|---|---|---|
| SP71-6163 VA vs. IACSP95-5000 VA | brown4 | 365 | 0.7 | 5 | 0.0084 |
| IACSP95-5000 NA vs. IACSP95-5000 AA | darkorange | 788 | 5.2 | 166 | 8.87E-198 |
| SP71-6163 AA vs. IACSP95-5000 AA | darkorange | 788 | 1.7 | 103 | 6.55E-156 |
| IACSP95-5000 AA vs. IACSP95-5000 VA | darkorange | 788 | 2.9 | 91 | 3.99E-106 |
| SP71-6163 NA vs. SP71-6163 AA | darkorange | 788 | 0.6 | 11 | 4.90E-10 |
| SP71-6163 NA vs. SP71-6163 VA | darkorange | 788 | 0.3 | 6 | 8.77E-06 |
| SP71-6163 NA vs. SP71-6163 VA | darkred | 1,147 | 0.4 | 4 | 0.0116 |
| SP71-6163 NA vs. IACSP95-5000 NA | ivory | 581 | 0.2 | 5 | 1.89E-05 |
| SP71-6163 VA vs. IACSP95-5000 VA | lightsteelblue1 | 814 | 1.5 | 11 | 7.05E-06 |
| IACSP95-5000 NA vs. IACSP95-5000 VA | lightsteelblue1 | 814 | 1.4 | 9 | 0.0002 |
| IACSP95-5000 AA vs. IACSP95-5000 VA | lightsteelblue1 | 814 | 3.0 | 10 | 0.0107 |
| IACSP95-5000 NA vs. IACSP95-5000 VA | magenta | 3,224 | 5.4 | 46 | 4.70E-28 |
| SP71-6163 VA vs. IACSP95-5000 VA | magenta | 3,224 | 5.7 | 47 | 1.25E-27 |
| IACSP95-5000 AA vs. IACSP95-5000 VA | magenta | 3,224 | 11.7 | 48 | 3.33E-15 |
| IACSP95-5000 NA vs. IACSP95-5000 VA | midnightblue | 1,637 | 2.7 | 10 | 0.0071 |
| SP71-6163 VA vs. IACSP95-5000 VA | midnightblue | 1,637 | 2.9 | 10 | 0.0092 |
| IACSP95-5000 NA vs. IACSP95-5000 AA | plum1 | 367 | 2.4 | 21 | 1.64E-12 |
| IACSP95-5000 AA vs. IACSP95-5000 VA | plum1 | 367 | 1.3 | 6 | 0.0227 |
| SP71-6163 NA vs. SP71-6163 AA | steelblue | 626 | 0.4 | 8 | 3.76E-07 |
| IACSP95-5000 NA vs. IACSP95-5000 AA | steelblue | 626 | 4.1 | 17 | 1.52E-05 |
| IACSP95-5000 NA vs. IACSP95-5000 AA | violet | 567 | 3.7 | 49 | 3.36E-37 |
| IACSP95-5000 AA vs. IACSP95-5000 VA | violet | 567 | 2.1 | 32 | 1.81E-26 |
| SP71-6163 NA vs. SP71-6163 AA | violet | 567 | 0.4 | 4 | 0.0112 |

NA: no aphids (control), AA: aviruliferous aphids, VA: viruliferous aphids.

**Table S9: Primer combinations used for amplifying and sequencing the genome of the sugarcane yellow leaf virus isolate employed in the experiments.**

| Primers | Forward / Reverse Sequences | Target region (nt) | $A_T$ | Reference |
|---|---|---|---|---|
| F1 / R13 | ACAAAATATATCGGGAGGGAAACCCCT / GTCCAATGGTTGTGTCTGGCAG | 1-340 | 60°C | Chinnaraja et al. (2013) / This study |
| F1 / R1 | ACAAAATATATCGGGAGGGAAACCCCT / GAGAGCCTCTTCCCCGTTYTTCAATC | 1-1,024 | 60°C | Chinnaraja et al. (2013) |
| F7 / R7 | TCATGATATATAGGACCGTCACG / GTTGAGCTGGTTGACTGGAGTGAA | 825-1,219 | 60°C | This study |
| R2 / R2 | CAYGTCACCACCRGGASACTCAGTGC / CCKGGGATRTCRCTGGTGGACTCTTC | 922-2,033 | 60°C | Chinnaraja et al. (2013) |
| F8 / R8 | CATCGTCTCGCAGTTGATTCAAAAG / GGTTCTACACGGTTCATATGCCTTG | 1,843-2,307 | 60°C | This study |
| F3 / R3 | GCAGCAGAACGGAGGGAAGAAGTC / TGAGTTTGGGCGTACARGACACCGCC | 1,933-3,041 | 60°C | Chinnaraja et al. (2013) |
| F9 / R9 | CTGGTAGATCAACTGGTAGCCCG / CGTGTGAGCAAAATTCTAGCTGTGA | 2,659-3,244 | 60°C | This study |
| F4 / R4 | CAACAACAACGAGCTAACCCGCCGCC / CTTGGCGTTCCTCTTGACGGGGAAGC | 2,940-4,088 | 57°C | Chinnaraja et al. (2013) |
| F10 / R10 | GGGATGTTCTCACTTTCACGGTTGA / GTTTCCTCTTGTTTTCACGATGTTG | 3,820-4,361 | 60°C | This study |
| F5 / R5 | CATGCTCCTCCGACGCAACTCCAGGTG / GGCTGGAGGCRTACTCTCCGTCCCTTC | 3,976-5,060 | 60°C | Chinnaraja et al. (2013) |
| F11 / R11 | CACTGATAAAATCCTGGAATGGGGTT / CCTCTCAAAGAACCACCGCTCA | 4,859-5,527 | 60°C | This study |
| F6 / ORF5 REV | GCTCGTTTRCCGACYCAGGTGAG / GCAGTGCCTCCCTGTATTCC | 4,962-5,880 | 60°C | Chinnaraja et al. (2013) / Abu Ahmad et al. (2006) |
| F12 / R12 | CTGTTCTTGGAGCCACTGACAC / CGCTAGCCCATCTGCATAGTTTTAG | 5,341-5,807 | 60°C | This study |
| F13 / ORF5 REV | TCTTCCTCCTCCTTGTTTGGTTGA / GCAGTGCCTCCCTGTATTCC | 5,655-5,880 | 60°C | This study / Chinnaraja et al. (2013) |

**Table S10: Information on sugarcane yellow leaf virus and potato leafroll virus isolates included in the phylogenetic analyses.**

| Accession code | Isolate | Country | Reference |
|---|---|---|---|
| PP796534 | BRA-SP1 | Brazil | This study |
| AF157029 | SCYLV-A | USA | Moonan et al. (2000) |
| AJ249447 | SCYLV-F | USA | Smith et al. (2000) |
| AM072750 | BRA-YL1 | Brazil | Abu Ahmad et al. (2006) |
| AM072751 | CHN-YL1 | China | Abu Ahmad et al. (2006) |
| AM072752 | PER-YL1a | Peru | Abu Ahmad et al. (2006) |
| AM072753 | PER-YL1b | Peru | Abu Ahmad et al. (2006) |
| AM072754 | REU-YL1a | Reunion | Abu Ahmad et al. (2006) |
| AM072755 | REU-YL1b | Reunion | Abu Ahmad et al. (2006) |
| AM072756 | REU-YL2 | Reunion | Abu Ahmad et al. (2006) |
| AY236971 | IND | India | Gaur et al. (2009) |
| GU190159 | CHN-FJ1 | China | Gao et al. (2012) |
| GU327735 | chn1 | China | Wang and Zhou (2010) |
| GU570006 | Haw87-4094 | USA | ElSayed et al. (2011) |
| GU570007 | Haw87-4319 | USA | ElSayed et al. (2011) |
| GU570008 | Haw73-6110 | USA | ElSayed et al. (2011) |
| HQ342888 | CHN-HN1 | China | Unpublished |
| JF925152 | IND1 | India | Chinnaraja et al. (2013) |
| JF925153 | IND2 | India | Chinnaraja et al. (2013) |
| JF925154 | IND3 | India | Chinnaraja et al. (2013) |
| JF925155 | IND4 | India | Chinnaraja et al. (2013) |
| KF477092 | GZ-GZ18 | China | Lin et al. (2014) |
| KF477093 | HN-CP502 | China | Lin et al. (2014) |
| KF680098 | CBLK97154 | India | Unpublished |
| KT960995 | Sorg1_1 | USA | ElSayed et al. (2018) |
| KT960996 | Sorg2_2 | USA | ElSayed et al. (2018) |
| KT960997 | Sorg3_3 | USA | ElSayed et al. (2018) |
| KY052165 | REU-YL11 | Reunion | Lu et al. (2021) |
| KY052166 | REU-YL15 | Reunion | Lu et al. (2021) |
| MF197921 | MU-AB110 | Mauritius | Unpublished |
| MF197922 | MU-AB193 | Mauritius | Unpublished |
| MF197923 | MU-AB194 | Mauritius | Unpublished |
| MF197924 | MU-AB197 | Mauritius | Unpublished |
| MF197925 | MU-SC1233 | Mauritius | Unpublished |
| MF622078 | COL-YL1a | Colombia | Unpublished |
| MF622079 | CUB-YL1a | Cuba | Unpublished |
| MN097766 | PI 157033 | USA | Boukari et al. (2021) |
| OL466944 | AR-01 | Argentina | Rott et al. (2023) |
| OL466945 | AR-02 | Argentina | Rott et al. (2023) |
| OL466946 | AU-01 | Australia | Rott et al. (2023) |
| OL466947 | BR-01 | Brazil | Rott et al. (2023) |
| OL466948 | BR-02 | Brazil | Rott et al. (2023) |
| OL466949 | BR-03a | Brazil | Rott et al. (2023) |
| OL466950 | BR-03b | Brazil | Rott et al. (2023) |
| OL466951 | BR-04 | Brazil | Rott et al. (2023) |
| OL466952 | BR-05 | Brazil | Rott et al. (2023) |
| OL466953 | BR-06-Leaf | Brazil | Rott et al. (2023) |
| OL466954 | BR-06-Sett | Brazil | Rott et al. (2023) |
| OL466955 | BR-07 | Brazil | Rott et al. (2023) |
| OL466956 | CN-01 | China | Rott et al. (2023) |
| OL466957 | CN-02 | China | Rott et al. (2023) |

| | | | |
|---|---|---|---|
| OL466958 | CO-01 | Colombia | Rott et al. (2023) |
| OL466959 | CO-02 | Colombia | Rott et al. (2023) |
| OL466960 | CO-03 | Colombia | Rott et al. (2023) |
| OL466961 | CO-04 | Colombia | Rott et al. (2023) |
| OL466962 | CO-05 | Colombia | Rott et al. (2023) |
| OL466963 | CO-06 | Colombia | Rott et al. (2023) |
| OL466964 | CO-07 | Colombia | Rott et al. (2023) |
| OL466965 | CO-08 | Colombia | Rott et al. (2023) |
| OL466966 | EC-01 | Ecuador | Rott et al. (2023) |
| OL466967 | GT-01a | Guatemala | Rott et al. (2023) |
| OL466968 | GT-01b | Guatemala | Rott et al. (2023) |
| OL466969 | GT-02a | Guatemala | Rott et al. (2023) |
| OL466970 | GT-02b | Guatemala | Rott et al. (2023) |
| OL466971 | GT-03 | Guatemala | Rott et al. (2023) |
| OL466972 | GT-04 | Guatemala | Rott et al. (2023) |
| OL466973 | GT-05 | Guatemala | Rott et al. (2023) |
| OL466974 | GT-06 | Guatemala | Rott et al. (2023) |
| OL466975 | GT-07 | Guatemala | Rott et al. (2023) |
| OL466976 | GT-08 | Guatemala | Rott et al. (2023) |
| OL466977 | GT-09 | Guatemala | Rott et al. (2023) |
| OL466978 | KR-01 | South Korea | Rott et al. (2023) |
| OL466979 | MU01-a | Mauritius | Rott et al. (2023) |
| OL466980 | MU01-b | Mauritius | Rott et al. (2023) |
| OL466981 | PG-01 | Papua New Guinea | Rott et al. (2023) |
| OL466982 | PG-02 | Papua New Guinea | Rott et al. (2023) |
| OL466983 | PH-01 | Philippines | Rott et al. (2023) |
| OL466984 | PH-02 | Philippines | Rott et al. (2023) |
| OL466985 | RE-01 | Reunion | Rott et al. (2023) |
| OL466986 | US-FL01a | USA | Rott et al. (2023) |
| OL466987 | US-FL01b | USA | Rott et al. (2023) |
| OL466988 | US-FL02 | USA | Rott et al. (2023) |
| OL466989 | US-FL03 | USA | Rott et al. (2023) |
| OL466990 | US-FL04 | USA | Rott et al. (2023) |
| OL466991 | US-FL05 | USA | Rott et al. (2023) |
| OL466992 | US-FL06 | USA | Rott et al. (2023) |
| OL466993 | US-HI01 | USA | Rott et al. (2023) |
| OL466994 | US-HI02 | USA | Rott et al. (2023) |
| OL466995 | US-LA01 | USA | Rott et al. (2023) |
| OL466996 | US-LA02 | USA | Rott et al. (2023) |
| OL466997 | US-LA03 | USA | Rott et al. (2023) |
| OL466998 | US-LA04 | USA | Rott et al. (2023) |
| OL466999 | US-LA05 | USA | Rott et al. (2023) |
| OL467000 | US-LA06 | USA | Rott et al. (2023) |
| OL467001 | US-LA07 | USA | Rott et al. (2023) |
| OL467002 | US-LA08 | USA | Rott et al. (2023) |
| OL467003 | US-LA09 | USA | Rott et al. (2023) |
| OL467004 | ZA-01a | South Africa | Rott et al. (2023) |
| OL467005 | ZA-01b | South Africa | Rott et al. (2023) |
| OL467006 | ZA-02 | South Africa | Rott et al. (2023) |
| OL467007 | ZA-03 | South Africa | Rott et al. (2023) |
| OL467008 | ZA-04 | South Africa | Rott et al. (2023) |
| NC_076505 | PLRV-C | Canada | Keese et al. (1990) |

**Table S11: Primer combinations used for RT-qPCR validation of the RNA-Seq libraries.**

| Forward / Reverse Sequences | Target | Product size (bp) | Reference |
|---|---|---|---|
| CACGGCCACTGGAAGCA / TCCTCAGGGTTCCTGATGCC | *GAPDH* | 101 | Iskandar et al. (2004) |
| GCTGTCACCCGGAACTACAG / AAGCGCTTTCAGTGTCGTCT | Cluster_58111 | 120 | This study |
| AGGATCCCAAGTTCACCGAC / CCGCTCTTTGAAGACGGACA | Cluster_3551 | 117 | This study |
| GAAGCCCTACCAAGCACACT / GTTTGTCAGGGCGGTCAGTA | Cluster_12386 | 100 | This study |
| TGGTGAACAAGACTCGCCTT / GCCAGCATATCCTCACTGCT | Cluster_41494 | 117 | This study |
| TCACCTCAGCTGAAGAACTCTC / GGCGGTCCCCCATATCATTT | Cluster_43698 | 119 | This study |
| GTGGCAACGGACTAACAGGA / CGCACACGGCACAACAATAA | Cluster_68973 | 111 | This study |
| GGTTACACCAGGACAGGAGC / GGGAAAAGGGGAAGTGGGAG | Cluster_75129 | 101 | This study |
| ATGTTGATGGCAGCAGCTCT / CGACATCCTTGCTCGCATTG | Cluster_81415 | 100 | This study |
| TGTGGAGAGGTTGTGGAAGA / CTCCCAAGCAAAAACGGCAA | Cluster_104101 | 113 | This study |
| GCTGTCACCCGGAACTACAG / AAGCGCTTTCAGTGTCGTCT | Cluster_58111 | 120 | This study |
| AGGATCCCAAGTTCACCGAC / CCGCTCTTTGAAGACGGACA | Cluster_3551 | 117 | This study |

### Supplementary Figures

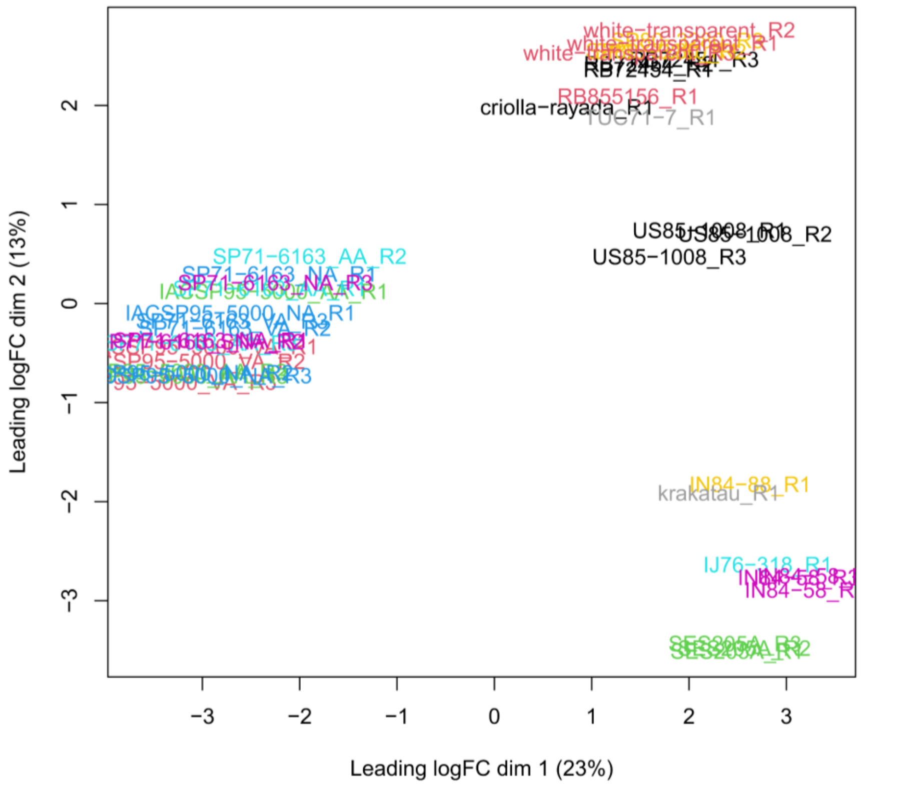


**Figure S1. Multidimensional scaling plot of RNA-Seq samples.** Genotypes included in the analyses are IN84-58, IN84-88, Krakatau, SES205A, IJ76-318, US85-1008, White Transparent, Criolla Rayada, TUC71-7, RB72454, SP80-3280, and RB855156, as well as SP71-6163 and IACSP95-5000 under three conditions: no aphids (NA, control), aviruliferous aphids (AA), and viruliferous aphids (VA).

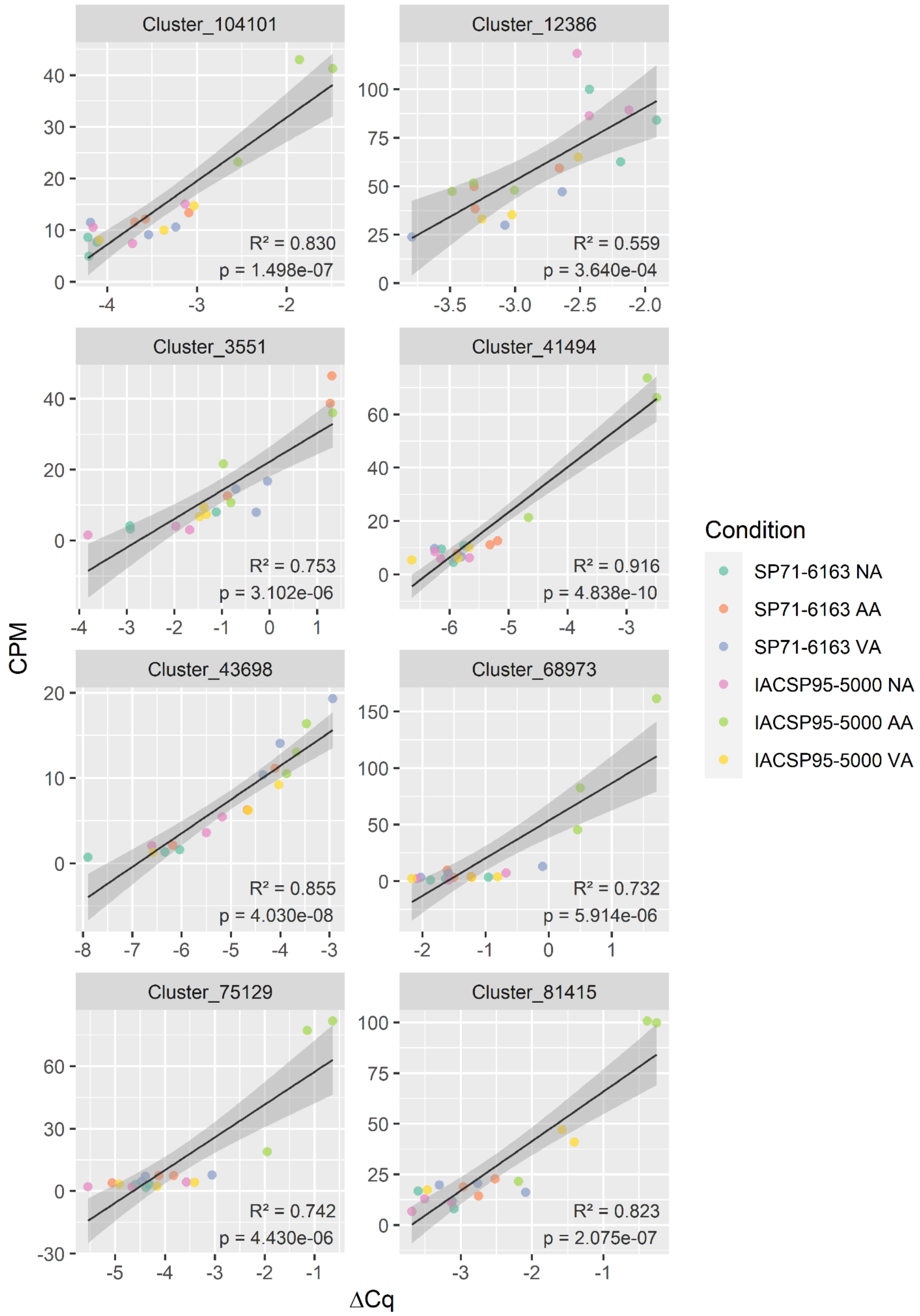


**Figure S2. Correlation plots between counts per million (CPM) values obtained by RNA-Seq analyses and quantitative cycle (ΔCq) values obtained by RT-qPCR for nine differentially expressed genes.** NA: no aphids (control), AA: aviruliferous aphids, VA: viruliferous aphids.

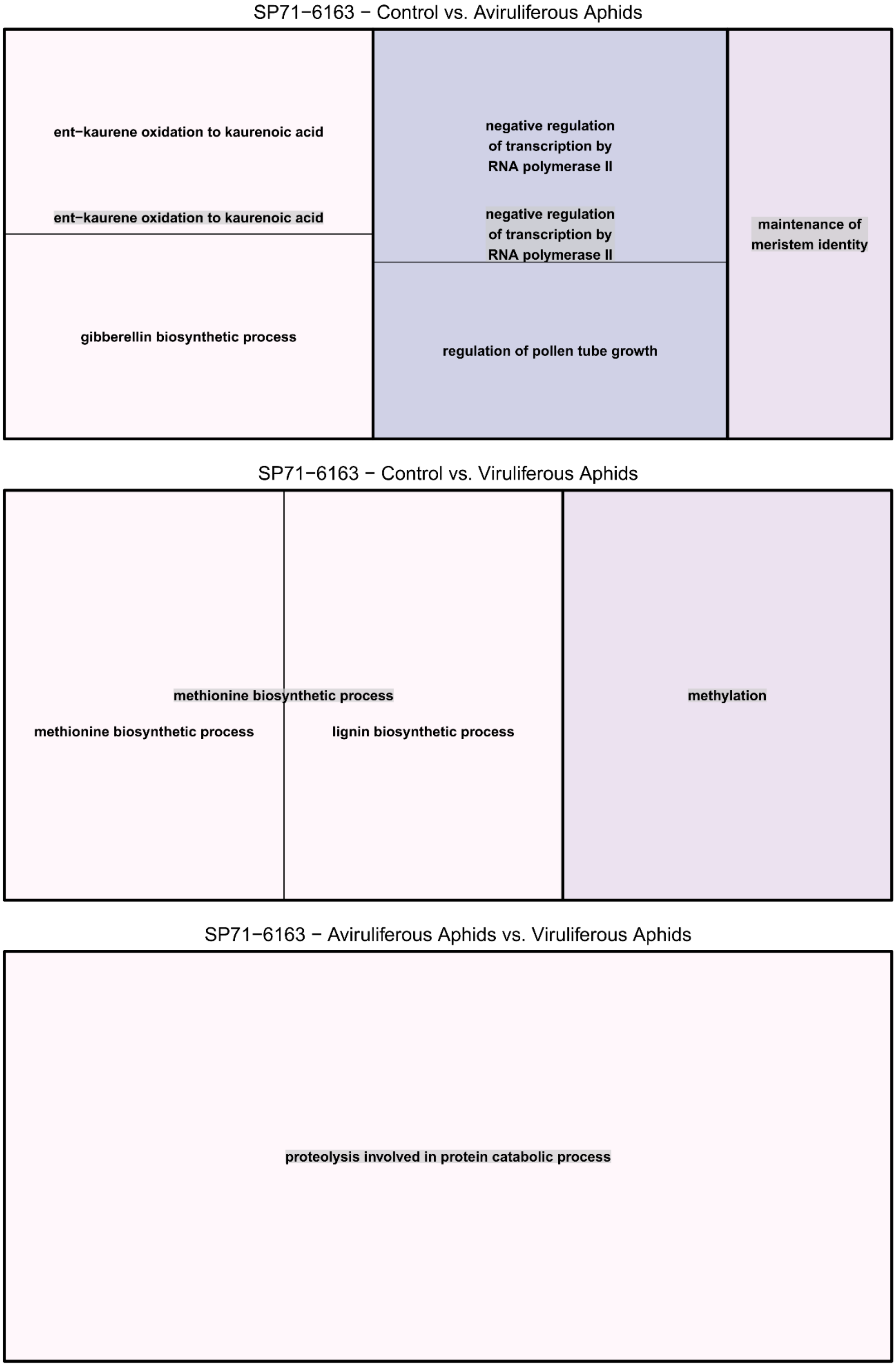


**Figure S3. REVIGO representation of Gene Ontology categories enriched in the sets of differentially expressed genes from comparisons of SP71-6163 under different conditions.**

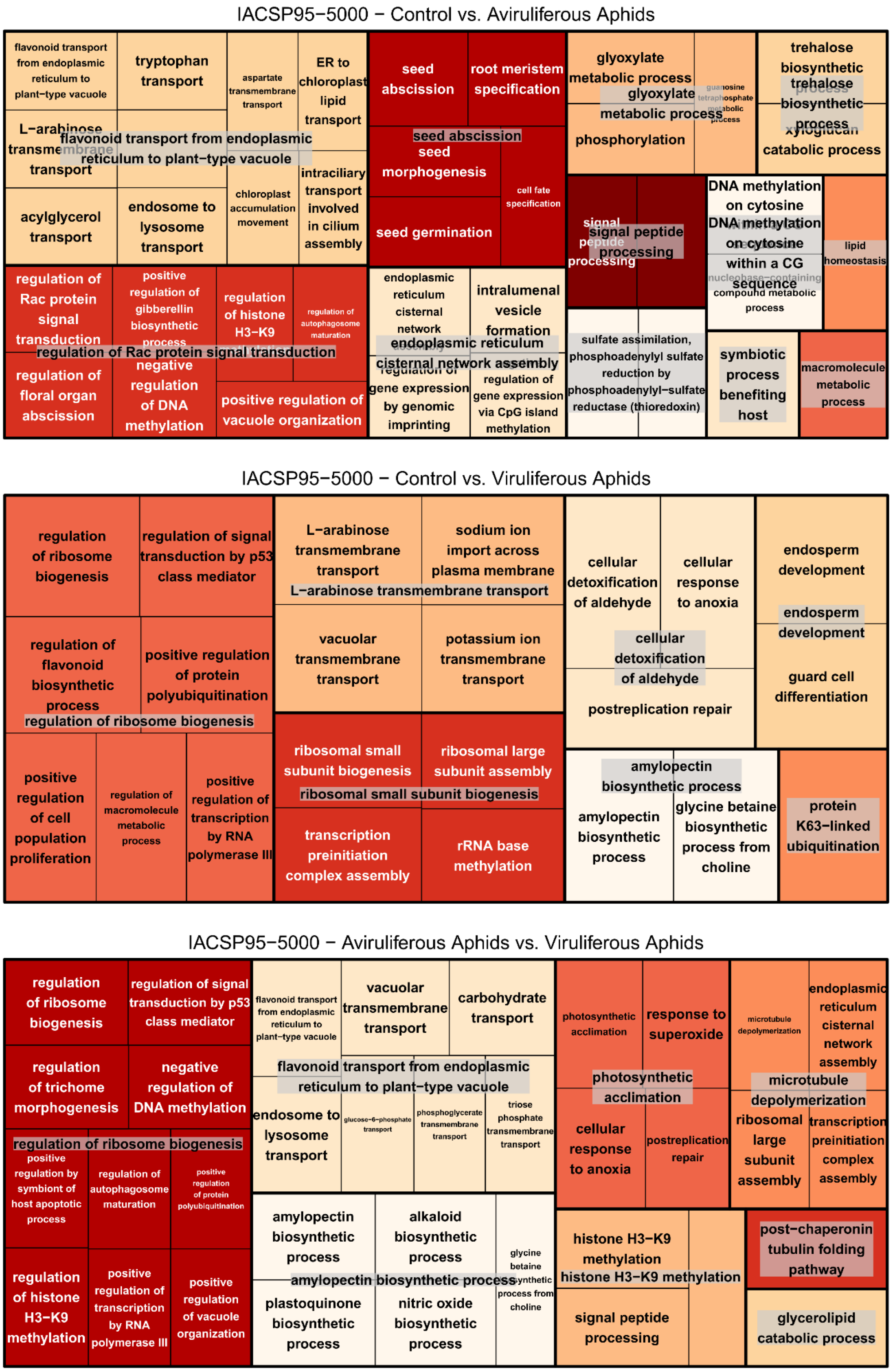


**Figure S4. REVIGO representation of Gene Ontology categories enriched in the sets of differentially expressed genes from comparisons of IACSP95-5000 under different conditions.**

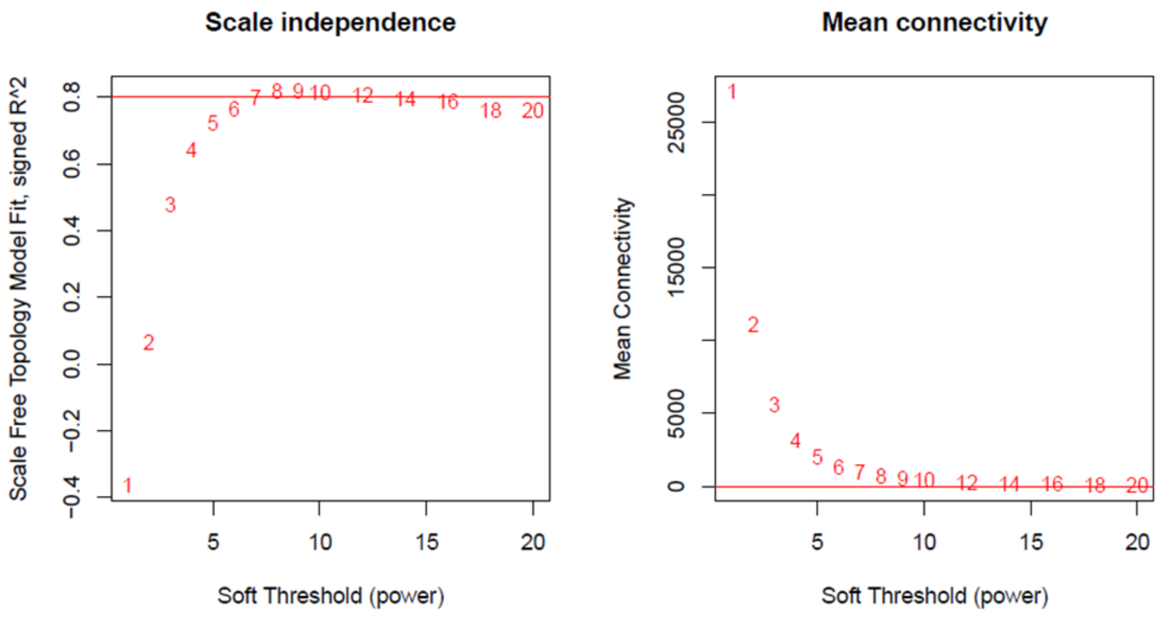


**Figure S5. Scale independence and average connectivity of the gene coexpression network according to the soft threshold value.**

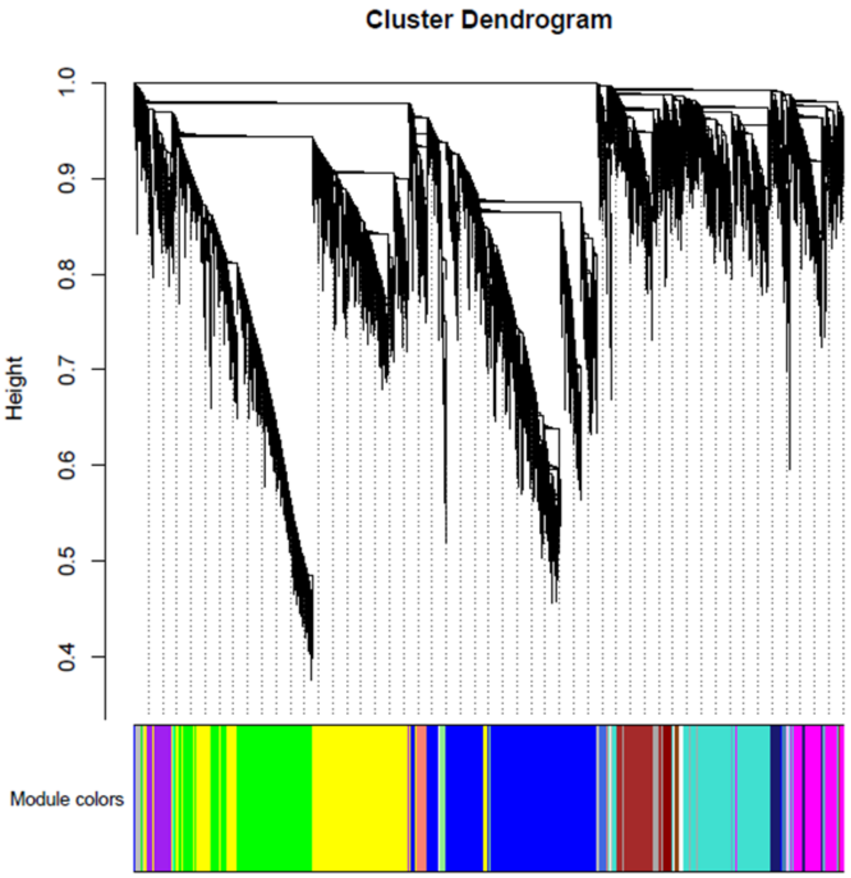


**Figure S6. Dendrogram of modules identified in the gene coexpression network.**

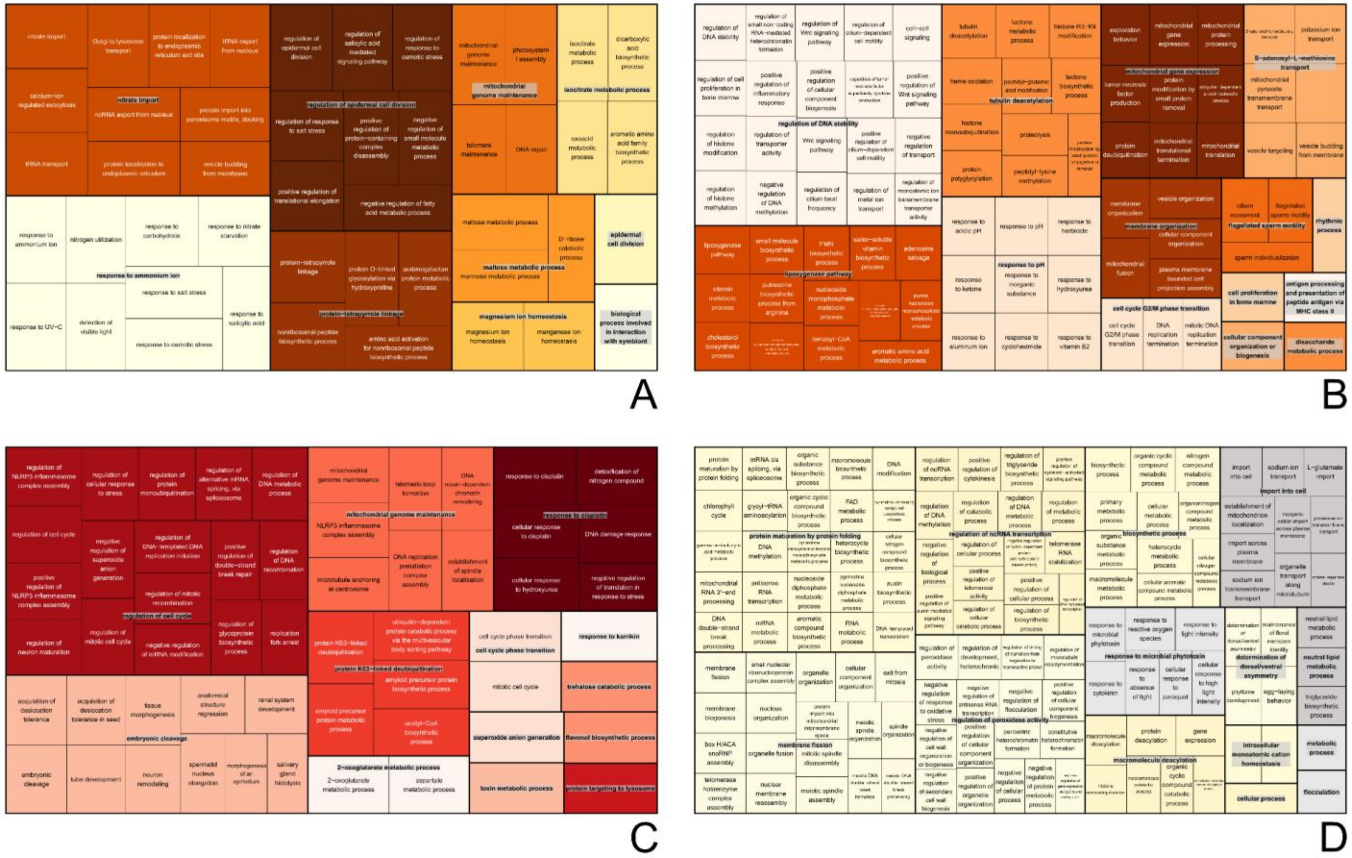


**Figure S7. REVIGO representation of Gene Ontology categories enriched in the network modules with overrepresentation of differentially expressed genes: (A) brown4, (B) darkorange, (C) darkred, (D) ivory, (E) lightsteelblue1, (F) magenta, (G) midnightblue, (H) plum1, (I) steelblue, and (J) violet.**

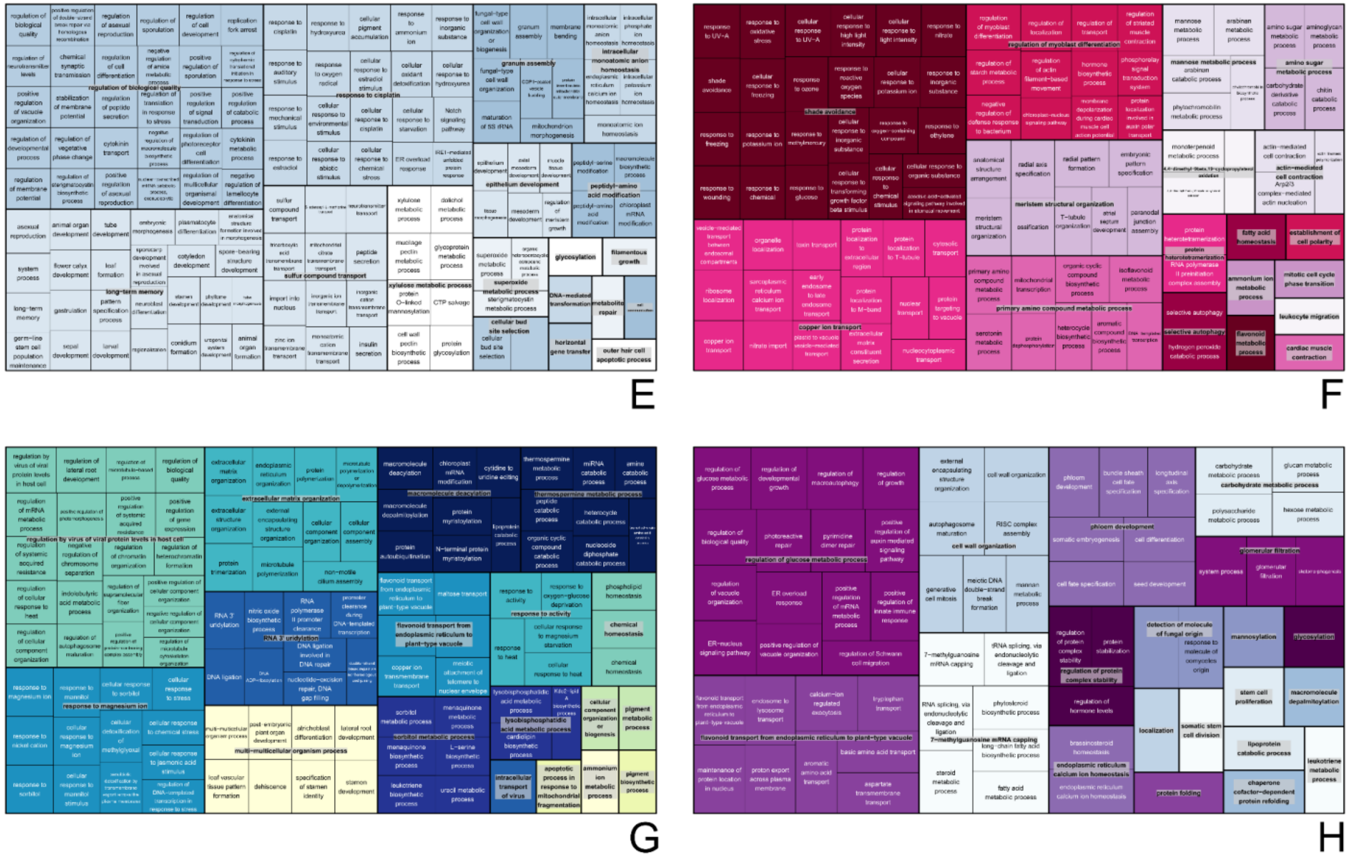


**Figure S7 (continued). REVIGO representation of Gene Ontology categories enriched in the network modules with overrepresentation of differentially expressed genes: (A) brown4, (B) darkorange, (C) darkred, (D) ivory, (E) lightsteelblue1, (F) magenta, (G) midnightblue, (H) plum1, (I) steelblue, and (J) violet.**

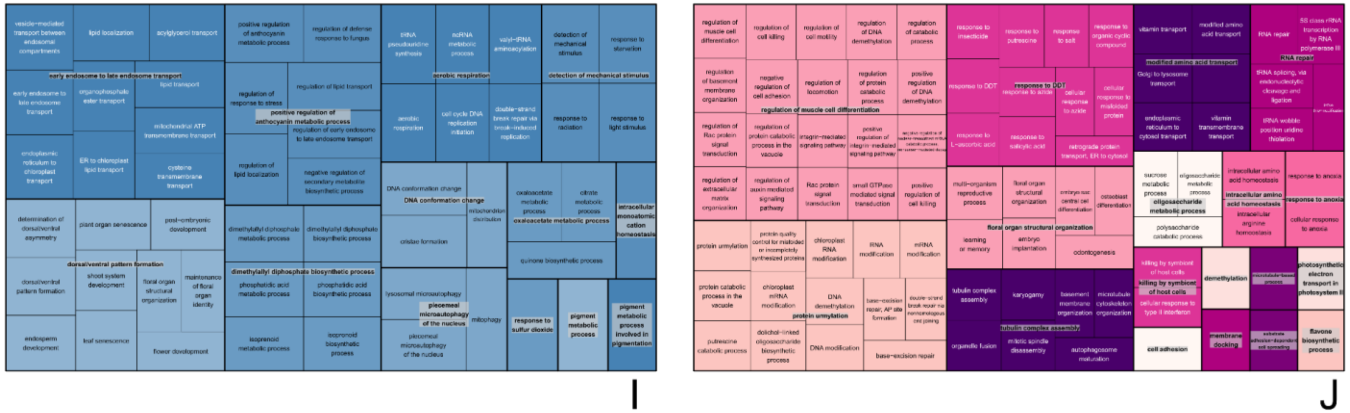


**Figure S7 (continued). REVIGO representation of Gene Ontology categories enriched in the network modules with overrepresentation of differentially expressed genes: (A) brown4, (B) darkorange, (C) darkred, (D) ivory, (E) lightsteelblue1, (F) magenta, (G) midnightblue, (H) plum1, (I) steelblue, and (J) violet.**
